# Learning to select computations in recurrent neural circuits

**DOI:** 10.64898/2026.04.14.718499

**Authors:** Sixing Chen, Frederick Callaway, Sreejan Kumar, Shira M. Lupkin, Joni D. Wallis, Vincent B. McGinty, Erin L. Rich, Marcelo G. Mattar

## Abstract

Two hallmarks of biological computation are its flexibility and efficiency. These features are often attributed to cognitive control processes that balance external utility against computational cost. However, how the brain could implement such adaptive control remains unknown. Here, we provide one possible answer by combining the computational theory of rational meta-reasoning with a meta-learning algorithm recently proposed as a model of prefrontal cortex. This yields a recurrent neural network model that learns to select computations. In simple choice tasks, the model approximates the algorithms and representations of optimal symbolic models and reproduces neural dynamics observed in macaque orbitofrontal cortex. In multi-step planning tasks, the model replicates key behavioral signatures of human planning strategies and captures human neural dynamics associated with step-by-step mental simulation. Our framework unifies meta-reasoning and meta-learning by showing that learning to reason can be understood as learning to learn from information generated by one’s own cognitive operations, providing a mechanistic account of how adaptive control of thought can be implemented in neural systems.

## 1 Introduction

Faced with the choice of whether to buy a bucket of popcorn at a movie theater, you might recall a previous experience of eating popcorn there, simulate the wait in line, or compare the price to what you paid last time. Each of these cognitive operations could influence your decision, yet each also takes time and effort. Humans and other highly intelligent animals routinely navigate such tradeoffs, deploying the right computations at the right time to solve complex problems despite severe limits on cognitive resources. This capacity for adaptively deciding what to think and when—known as “meta-reasoning”—is central to intelligent behavior (Gershman et al., 2015; Griffiths et al., 2019; c.f. Howes et al., 2009; Shenhav et al., 2013).^1^ Understanding how humans flexibly regulate their own thought processes is thus essential to explaining human intelligence and building human-like artificial agents.

Previous research has proposed that human thought processes are resource-rational, that is, optimized to balance the external utility of one’s actions with the cost of the computations used to select those actions. This perspective has led to significant progress in uncovering human cognitive strategies in various domains (Lieder and Griffiths, 2017; Lieder et al., 2018; Callaway et al., 2021, 2022, 2024a). However, the approach faces two major challenges.

The first challenge is algorithmic: identifying optimal computational strategies is itself a costly computational process—so costly, in fact, that a founder of rational meta-reasoning ultimately abandoned it (Russell, 1997). To circumvent this problem, previous approaches have either ignored cognitive and neural plausibility (Mattar and Daw, 2018; Callaway et al., 2022) or optimized over a small set of researcher-specified strategies (Howes et al., 2009; Lieder and Griffiths, 2017). While some work has proposed that computational strategies could be learned (Callaway et al., 2017; Lieder et al., 2018; He and Lieder, 2025), concrete implementations have relied on domain-specific, hand-engineered features. This leaves open the question of how a biological agent could ever acquire even a near-optimal strategy for selecting computations.

The second challenge is representational. A computational strategy is defined by both an algorithm and the representations over which that algorithm operates. Nevertheless, most previous research has focused exclusively on the algorithm, while assuming a symbolic representational space based on cognitive architectures (Howes et al., 2009), classical AI methods (Mattar and Daw, 2018; Callaway et al., 2022), or Bayesian inference (Drugowitsch et al., 2012; Callaway et al., 2021). It thus remains unclear how meta-reasoning could operate in a distributed neural system such as the brain, and even less clear how one could find evidence for such a process in neural activity.

A path forward follows from a key insight: the problem of selecting computations is analogous to the problem of selecting actions, and both can be formalized using the tools of statistical decision theory (Matheson, 1968; Russell and Wefald, 1991; Hay et al., 2012). In particular, computations can be modeled as internal actions that generate decision-relevant information to reduce uncertainty about which external action is best. In the popcorn example above, recalling a past experience of eating popcorn or simulating the wait in line both provide information about whether you should buy a bucket. Computations thus function as internal “experiments” whose results update the agent’s beliefs about the decision at hand. Accordingly, controlling a sequence of such computations is directly analogous to controlling a *learning* process. This suggests that a neurally plausible theory of computation selection could be built on a neurally plausible theory of learning.

However, the timescale of thought processes poses a critical constraint. Learning driven by slow synaptic plasticity cannot explain the speed and flexibility of human thought. Instead, meta-reasoning requires a system that learns quickly from few samples and generalizes across a wide range of problems. One prominent theory proposes that such fast learning could arise from meta-reinforcement learning (meta-RL), in which fast adaptations are implemented by recurrent dynamics in prefrontal cortex (PFC), which are themselves learned through slowly adjusting synaptic weights across tasks (Wang et al., 2018). Yet most meta-RL models focus exclusively on interactions with the external environment. Recent work has begun to extend this framework to include internal computations (Jensen et al., 2024), but still only optimizing over *how much* computation to perform, falling short of explaining the richness and flexibility of human mental strategies. Moreover, the neural mechanisms underlying computation selection remain only minimally explored. This leaves a critical gap in understanding whether—and if so, how—adaptive control of computations can be supported by neural dynamics in PFC.

Here, we show that a meta-RL agent augmented with mental actions can learn adaptive computational strategies that approximate rational meta-reasoning. The agent learns to select not only when to perform computations but also which computations to perform, with each computation generating information and updating its internal state. In a simple choice task, the agent learns strategies consistent with optimal symbolic models and reproduces neural dynamics observed in macaque orbitofrontal cortex (OFC). In a more complex planning task, the agent learns a strategy resembling best-first search and reproduces key patterns in human eye-tracking data. Further, the agent captures neural dynamics associated with step-by-step mental simulation found in human hippocampus and PFC. Our work establishes a framework that bridges normative theories of optimal computation selection with underlying neural mechanisms, providing a principled theory for how meta-reasoning can be implemented in neural systems.

## 2 Results

### 2.1 Meta-learning to select computations

To model the rapid and flexible selection of computations, we develop a recurrent neural network (RNN) trained with meta-RL (Figure 1). The model uses gated recurrent units (GRUs) with an actor–critic architecture. The recurrent dynamics evolve as

**Figure 1:**
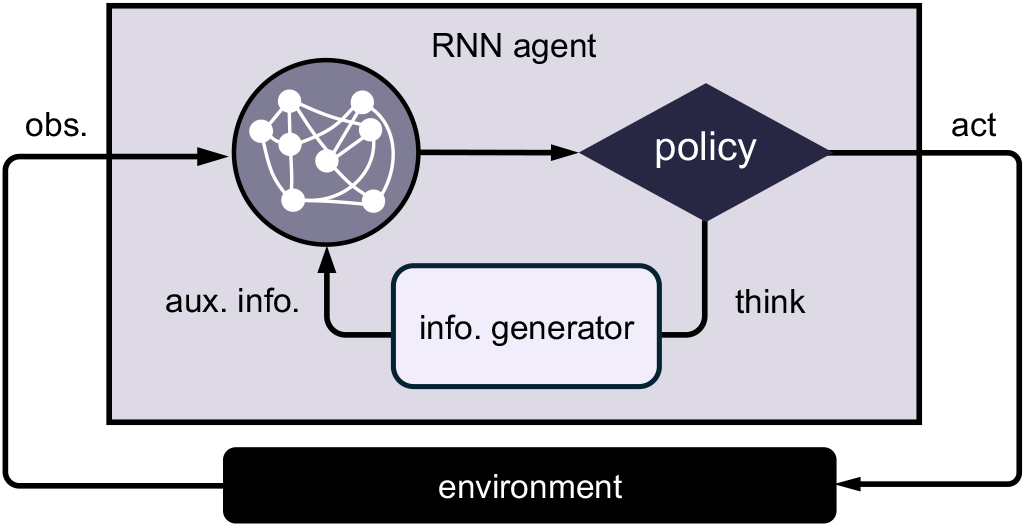
Model schematic. The agent consists of an RNN that receives observations from the external environment and selects actions. The actions can be either physical (“act”) or mental (“think”) actions. Physical actions alter the environmental state. Mental actions leave the environmental state unchanged and instead query an information generator, which returns decision-relevant information that is appended to the agent’s input at the subsequent time step.

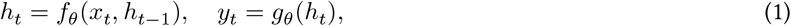

where *θ* denotes the RNN parameters, *h*_*t*_ is the hidden state, *x*_*t*_ is the input, and *y*_*t*_ is the output. The output consists of an action policy *π*_*θ*_(*a*_*t*_ | *h*_*t*_) over valid actions and a value baseline *V*_*θ*_(*h*_*t*_). Crucially, the agent’s action space includes not only “physical” actions (buying popcorn), but also computational, or “mental,” actions (wondering how it tastes). Unlike physical actions, which alter the environmental state, mental actions leave the environment unchanged and instead query an “information generator”—a module that implements task-specific cognitive operations such as recalling a memory or simulating a future action. The generator returns decision-relevant information that is included in the next input (*x*_*t*+1_) so that it can be integrated into the agent’s hidden state. Each mental action thus provides information but incurs a cost (a negative reward). The reasoning process corresponds to sequentially selecting mental actions that refine the agent’s internal “mental” state, trading off the value of additional information against the computational cost of producing it. As is standard in meta-RL, we additionally provide as inputs the previous action *a*_*t*−1_ and the previous reward *r*_*t*−1_ (Wang et al., 2016, 2018; Duan et al., 2016). The agent is trained with policy gradient methods to maximize the expected cumulative reward:

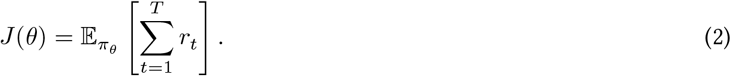

Importantly, the rewards here include both computational cost (for mental actions) and external utility (for physical actions). After training, the agent’s parameters are frozen during evaluation, with all adaptation achieved only by its recurrent dynamics.

This architecture provides a concrete implementation of rational meta-reasoning in neural systems. Classically, rational meta-reasoning formalizes computation selection as a “metalevel Markov decision process” (metalevel MDP), where an agent sequentially selects costly computations to update its belief state and guide behavior (Russell and Wefald, 1991; Hay et al., 2012). The optimal policy for a metalevel MDP is a computational strategy that maximizes expected external utility minus internal computational costs. Our architecture generalizes this formalism by training the agent with the same objective as the metalevel MDP while letting the effect of computation (and, therefore, the representational content of the internal state) be learned in the recurrent dynamics rather than hand-specified.^2^

We interpret this architecture at two levels. Cognitively, we interpret the RNN as a central controller that coordinates one or more cognitive modules (e.g. a world model or a memory system) to guide behavior. The information generator is an abstraction of those modules: given a query from the controller (e.g., “how expensive was the popcorn last time?”), it returns a relevant bit of information (e.g., a reinstated experience of feeling ripped off). Biologically, we interpret the execution of mental actions as interactions between PFC (the RNN) and other brain regions, such as hippocampus, basal ganglia, amygdala, and cerebellum (the information generator). These regions have been proposed to read out task-relevant information from PFC activity, carry out specialized computations (e.g., mental simulation, working memory gating, or episodic recall), and feed the results back to update PFC representations (Jensen et al., 2024; O’Reilly and Frank, 2006; Wassum, 2022; Stopper et al., 2014; Pemberton et al., 2024). In this view, the querying process corresponds to interactions between brain regions that support flexible sequential computation.

We next test whether this model learns effective computation selection strategies and reproduces behavioral and neural signatures observed during deliberation.

### 2.2 Meta-RL agent learns to select computations on a simple choice task

We first tested whether the meta-RL agent can learn to select computations in a simple choice task where human behavior has been well characterized by an optimal symbolic model (Callaway et al., 2021). In this task, participants’ gaze was recorded as they chose between pairs or triplets of snack foods. They could freely view the items before selecting the one they would rather receive at the end of the study (Figure 2A-B). Consistent with the literature on attention and decision-making (Orquin and Loose, 2013), we interpret the gaze behavior primarily as an indicator of the underlying computational process rather than sensory information gathering (as justification: patterns of gaze and choice are similar when the items are bound to locations on the screen but not actually visible; Weilbächer et al., 2021; Ferro et al., 2024). Eye gazes thus offer an observable proxy for otherwise unobservable deliberation.

**Figure 2:**
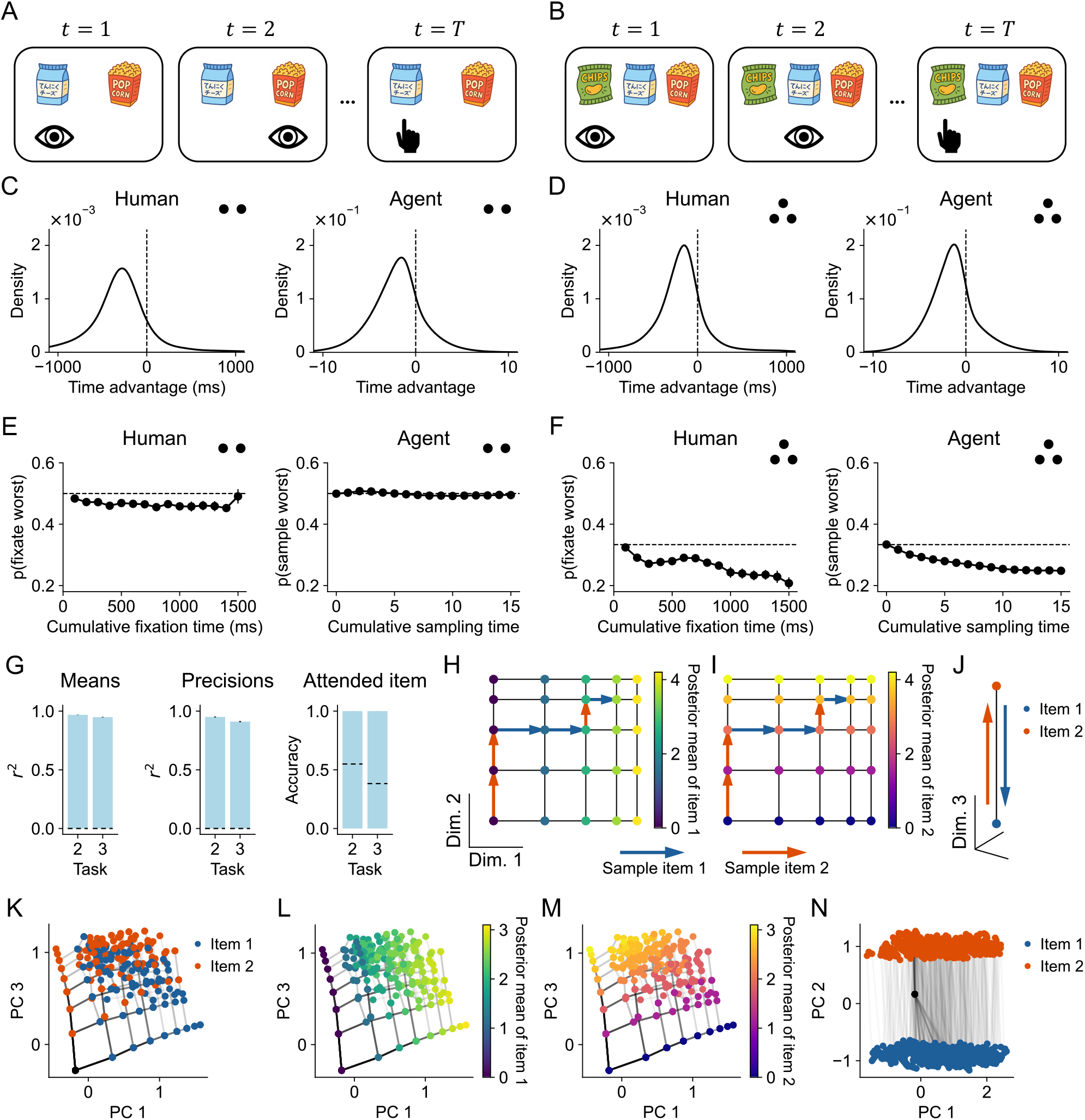
Meta-RL agent learns to select computations on a simple choice task. (A–B) Illustration of binary (A) and trinary (B) choice tasks. Human participants freely view the items and then select the one they would rather eat. (C–D) Distribution of time advantage, defined as the cumulative time attending to the current item minus the mean cumulative time attending to the other item(s) from trial onset to the current time. (E–F) Probability of attending to the worst item as a function of the cumulative time since the start of the trial. Dashed lines show chance levels. Number of dots indicates binary or trinary choice tasks. (G) Decoding performance for belief-state sufficient statistics. Dashed lines show baselines computed by randomly shuffling labels. (H–J) Schematics of belief state representation. Dots indicate hidden states and arrows indicate transitions driven by sampling actions. Dimension 1 and 2 encodes posterior means of item 1 and 2, respectively. Dimension 3 encodes the current attended item. Hidden states are colored by posterior mean of item 1 (H), item 2 (I), and the current attended item (J). Sampling an item moves the hidden state along the corresponding dimension. (K–N) PCA results. Panels K to M show an example condition in which both items have a value of 4. Hidden states are colored by posterior mean of item 1 (L), item 2 (M), and attended item (K and N). Line transparency shows transition frequency. Panels K to N are simulated with noise-free value samples for visualization (Methods). Error bars show standard errors across participants or random seeds.

Following Callaway et al. (2021) (c.f., Tajima et al., 2016; Jang et al., 2021), we formalize the setting as a sequential sampling problem. At each time step, the agent could attend to one item. Attending to an item queries the information generator, which returns a noisy sample centered on the item’s true value; this provides information but incurs a cost. The input to the agent included the identity of the currently attended item, the value sample from that item, the previous action, and the previous reward. The action space comprised sampling actions (attend to an item) and decision actions (commit to an item), with 2 or 3 of each for the binary and trinary tasks, respectively. Each action, whether sampling or decision, maps to a specific item. Each sampling step incurs a cost, with an additional switching cost if the currently sampled item is different from the previous one. The agent thus faces the problem of deciding whether to make a decision or to continue sampling, and if sampling, which item to sample from.

We tested whether the agent learned two key features of the optimal policy: (i) the tendency to sample items with more uncertain value estimates, and (ii) the tendency to focus on items with the highest and second-highest value estimates (Callaway et al., 2021). The rationale is straightforward: sampling is only useful if it has the potential to change the decision policy. This is most likely when either (i) the value estimate of an item is highly uncertain and therefore easily shifted, or (ii) the value estimate is close to its current competitor (the second-best item for the best item and the best item for all other items). To test the first feature, we computed the distribution of relative cumulative fixation time (“time advantage”) at the beginning of each new fixation, excluding the first fixation. For both humans and the agent, the distributions were centered below zero, indicating a tendency to fixate items that had received less attention and therefore had more uncertain value estimates (Figure 2C-D). To test the second feature, we examined the probability of sampling the worst item as a function of the cumulative sampling time. Humans initially viewed the worst item at chance levels, but in the three-item task, this probability quickly declined as attention shifted toward the top two items (Figure 2E-F). The agent exhibited the same decline. Indeed, the agent reproduced most of the key patterns in human behavior captured by the optimal symbolic model (Callaway et al., 2021; Figure S1), confirming that it had learned to select computations strategically.

Having established that the agent’s behavior matches the optimal strategy, we next asked whether it also learned to represent and update Bayesian belief states as in the optimal symbolic model. Specifically, the symbolic model maintains Gaussian posteriors over each item’s value given all previous value samples; this belief can be represented by the posterior mean and precision for each item. The model also tracks the identity of the currently attended item, since this determines which item can be sampled without paying the switch cost. To test for such representations, we trained linear decoders to decode these sufficient statistics from the agent’s hidden states. We found that all variables could be decoded with near-perfect accuracy in both tasks (Figure 2G), suggesting that the agent learned to represent belief states.

To understand the geometric structure of these representations, we applied principal component analysis (PCA) to the agent’s hidden states in the binary choice task. We hypothesized that the posterior means of the two items are encoded along two distinct dimensions (Figure 2H-I), with a third dimension encoding the identity of the attended item (Figure 2J). Sampling an item shifts the hidden state along the corresponding dimension. This yields two predictions. First, the hidden states’ projections in the posterior-mean subspace should be invariant to sample order. For example, assuming the value samples are the same, sampling left–then–right should land in the same position in the subspace as right–then–left, producing a grid-like structure (Figure 2I). Second, the step size induced by each sampling step in this subspace should shrink with more samples, since posterior means become increasingly precise. Our analyses confirmed both predictions: two principal components spanned a subspace where each component corresponded to the posterior mean of an item, the representation showed a certain degree of invariance to sample order, and step size decreased with more sampling (Figure 2K-M; Figure S2). Moreover, we identified a third principal component, orthogonal to this subspace, that encoded the identity of the currently attended item (Figure 2N). These findings demonstrate that the agent encodes the sufficient statistics of belief states in distinct subspaces and updates them dynamically through its hidden dynamics.

Together, these results provide a comprehensive picture of how the meta-RL agent learns to select computations in a manner consistent with the optimal symbolic model: the agent encodes and updates belief states in its hidden representations (Ortega et al., 2019; Mikulik et al., 2020; Alver and Precup, 2021), and uses its policy head to map belief states onto the learned policy. This establishes a direct connection between the normative framework of meta-reasoning and our neural network model of computation selection.

### 2.3 Meta-RL agent reproduces neural dynamics in macaque orbitofrontal cortex

Our framework predicts that computation selection is reflected not only in behavior (e.g., eye fixations) but also in neural dynamics in PFC. To test this prediction, we turned to a study by Rich and Wallis (2016), in which macaques first learned associations between pictures and rewards and were then trained on a binary choice task to select the higher-valued picture. During deliberation and choice, neural activity in the orbitofrontal cortex (OFC) was recorded. The authors applied linear discriminant analysis (LDA) to classify population activity moment-by-moment into subjective value categories (ranging from 1 to 4). The decoder’s output probabilities revealed that OFC representations alternated between the value of the chosen and unchosen options, indicating a deliberation process in which each option was considered in turn (Figure 3A).

**Figure 3:**
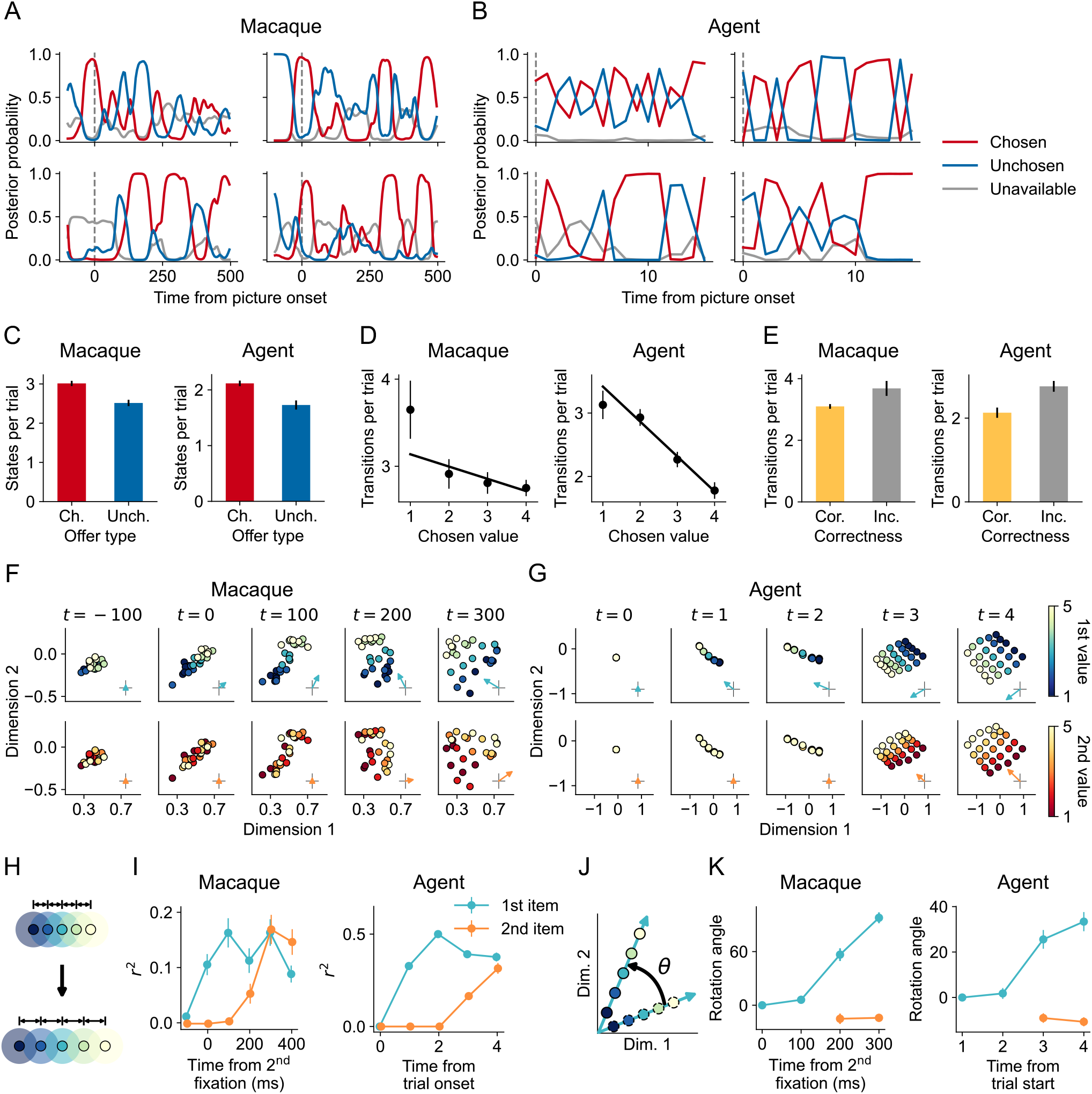
Meta-RL agent reproduces neural dynamics in macaque orbitofrontal cortex. (A–B) Posterior probabilities from the LDA decoder for chosen (red), unchosen (blue) and unavailable options (gray, average of both unavailable options), shown for 4 example trials. (C) Number of decoded value states for chosen and unchosen options. (D) Number of transitions between decoded states as a function of chosen value. (E) Number of transitions in correct versus error trials. (F–G) Temporal evolution of value-subspace representation in macaques (F) and the agent (G). Dimensions 1 and 2 are orthogonal bases within the value subspace. Each dot shows average neural activity for a given pair of values of the first- and second-viewed options. The first and second rows are colored by the value of the first- and second-viewed options, respectively. Arrows indicate the value gradient’s direction and magnitude. Macaque neural activity is aligned to the onset of the second fixation (approximately 200 ms after the first fixation); agent hidden activity is aligned to trial onset. Panel (F) shows an example session, and panel (G) shows an example agent. (H) Schematic of value gradient expansion. Decodability increases as neural activity between values become more separable. (I) Decoding performance of option value from the value subspace over time. (J) Schematic of value gradient rotation *θ* indicates rotation angle. (K) Rotation angle of the value gradient over time. Rotation angle is baseline-corrected relative to first-viewed option’s value gradient at *t* = 0 ms for macaques and *t* = 1 for the agent. Error bars show standard errors across sessions or random seeds.

We asked whether the meta-RL agent, trained on this task, would reproduce the same alternation. The task closely paralleled the earlier simple choice task but with one modification: the agent was required to sample for a minimum number of time steps (Methods). This mirrored the experimental constraint in Rich and Wallis (2016) where macaques had to maintain fixation on an option for a minimum amount of time to select it. And because this requirement constrained macaques’ eye movements, which could have the effect of decoupling deliberation from fixations, gaze could no longer serve as a readout of internal deliberation. We applied LDA analysis to the agent’s hidden states using an analogous procedure (Methods). The decoder’s output probabilities showed alternation patterns that reproduced the findings of the original study (Figure 3B). Importantly, the value states identified in the agent’s hidden dynamics exhibited statistical patterns closely resembling those in macaque OFC. In both cases, there were more value states corresponding to the chosen option than to the unchosen one (macaques: *t*(43) = 5.23, *p <* .001; agents: *t*(4) = 9.35, *p <* .001; Figure 3C). Moreover, higher value of the chosen option predicted fewer transitions between value states (macaques: *β* = −0.21, *p <* .001; agents: *β* = −0.62, *p <* .001; Figure 3D), whereas higher value of the unchosen option value predicted more transitions (macaques: *β* = 0.32, *p <* .001; agents: *β* = 0.18, *p <* .001). There were more transitions in error trials than in correct trials (macaques: *t*(42) = 2.77, *p* = 0.008; agents: *t*(4) = 11.89, *p <* .001; Figure 3E). These results suggest that the alternations observed in both macaque OFC and the agent’s hidden dynamics are not arbitrary fluctuations. Instead, they reflect the underlying computation selection process during deliberation.

To further understand how computations shape neural representations, we reanalyzed another study where macaques similarly learned to choose between two options (McGinty and Lupkin, 2023). In this task, the authors used visual crowding to prevent evaluation via peripheral vision, and macaques reported decisions with a manual level press instead of fixations. This design effectively couples fixations with deliberations. The authors reported that OFC activity encoded value estimates in a two-dimensional subspace. We reanalyzed the dataset to trace how this subspace representation unfolded over time. Following the authors’ approach, we aligned neural activity to the start of the second fixation, which was around 200 ms after the first fixation. We trained linear decoders to decode each option’s value using spike activity during a period when both values had relatively stable representations (200 to 400 ms after the second fixation). The corresponding regression weights spanned the two-dimensional value subspace. To examine how the value representations evolved over time, we then identified two orthogonal bases within the value subspace and projected spikes from different time points onto this subspace (Methods). The analysis revealed that the value gradients emerged following the order of fixations: the value gradient of the first-viewed option across trials emerged earlier (Figure 3F, first row), followed by that of the second (Figure 3F, second row). Correspondingly, the value of the first option became decodable in the value subspace before that of the second (Figure 3H-I).

We then tested whether the agent exhibited the same sequential emergence of value representations. We trained the agent on this task and, during evaluation, had it sample each option for two time steps to ensure equal coverage across value pairs and enable temporal alignment (Methods). Because sampling actions influenced the agent’s hidden state immediately, we aligned hidden activity to trial onset. Consequently, we expected the value gradient to emerge after the first time step (*t* = 1) for the first option and two time steps after (*t* = 3) for the second. Applying the same value-subspace analysis as in the macaque data revealed similar dynamics in the agent: the value gradient of the first-viewed option emerged after sampling the first option (Figure 3G, first row), and that of the second-viewed option emerged after sampling the second (Figure 3G, second row), with option values becoming decodable immediately after being sampled (Figure 3I). The close correspondence between macaques and the agent indicates that OFC neural geometry reflects belief-state representational geometry shaped by individual computations, providing mechanistic evidence that OFC supports the adaptive control of computations.

Intriguingly, we observed not only the emergence of value gradients but also rotation of these gradients over time (Figure 3J). As information about the second-viewed option came in, the gradient of the first-viewed option did not simply persist in place, but rotated to a new direction (Figure 3K). Simultaneously, the gradient of the second-viewed option emerged along a direction closely aligned with the original gradient of the first-viewed option. This suggests that both the macaque and the agent represent values with temporally organized subspaces, consistent with recent proposals that PFC employs temporal subspaces (or “structured slots”) to represent sequential information (Xie et al., 2022; Whittington et al., 2025).

### 2.4 Meta-RL agent learns human-like planning strategies on a planning task

Having validated the framework in a simple choice task where the optimal strategy has been well characterized, we next turned to a planning setting with a complex state space. In a task developed by Callaway et al. (2024b), participants navigated a tree-structured directed graph, aiming to maximize the sum of rewards collected at each state they visited (Figure 4A). On each trial, participants were presented with a new graph and had 15 seconds to identify and execute a plan, clicking on a sequence of states to visit them and collect their rewards (Figure 4B). At each step, they could only click on one of the two states that were connected to their current state. The trial ended when they reached a state with no outgoing connections or 15 seconds had elapsed; in the latter case, random moves were forced until a terminal state was reached. Finally, the authors adopted a gaze-contingent display, such that a state’s reward was only visible when the state was fixated. This ensured that the participant’s gaze was highly indicative of the state they were currently attending to (Figure 4C).

**Figure 4:**
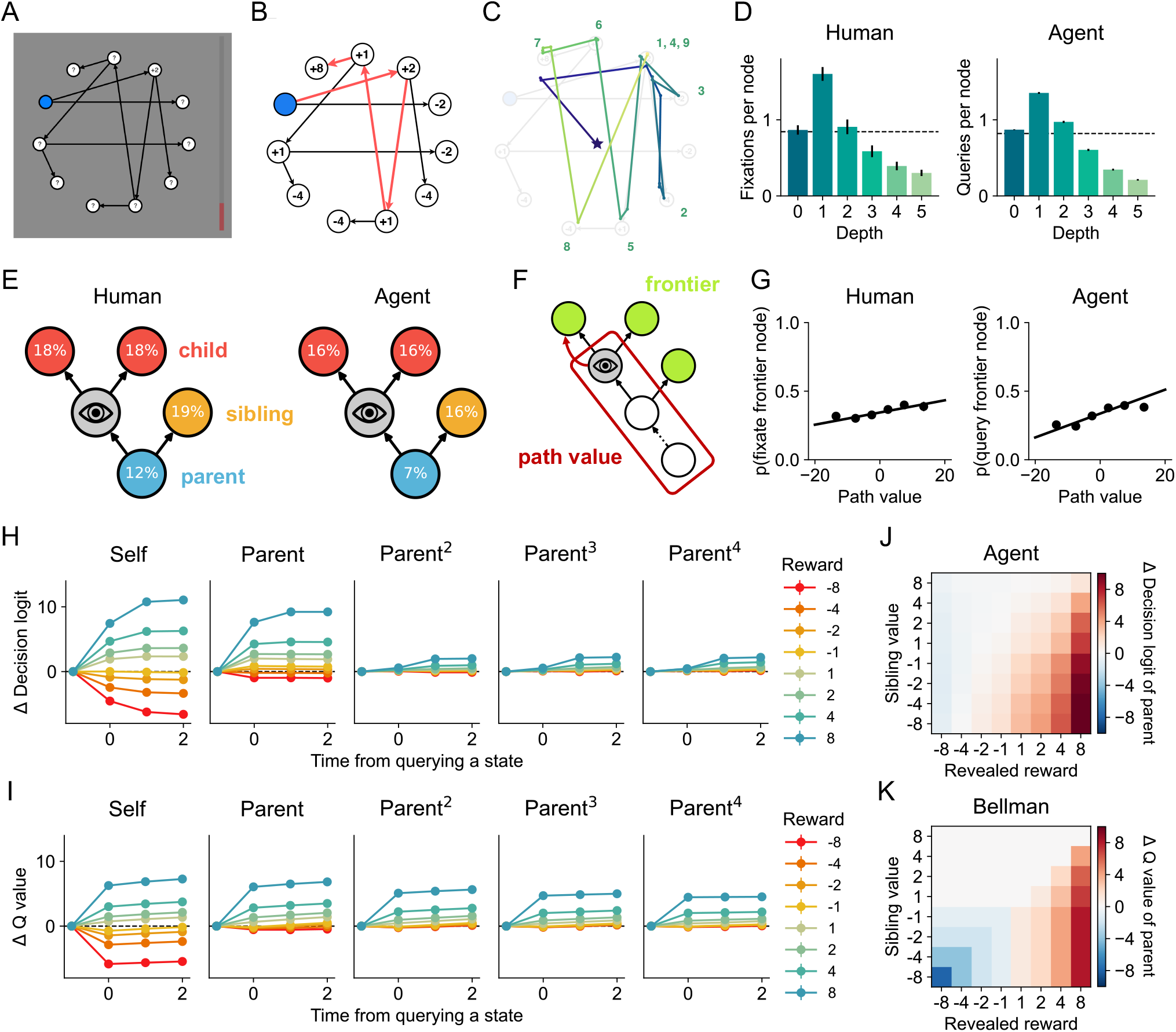
Meta-RL agent learns a human-like planning strategy. (A) Task interface. A visually-displayed decision tree is shown where eleven states are arranged on a circle, each associated with a certain reward. The current state is highlighted in blue and possible actions are indicated by arrows. Participants aim to select a path that maximizes cumulative reward by clicking on states. Each state’s reward is only revealed while the state is fixated. The bar on the right indicates the time left for the trial. (B) An example chosen path indicated by red arrows. (C) An example fixation sequence prior to the first action of the trial. Numbers indicate the order of fixations. (D) The average number of fixations or queries per state at different depths. Dashed lines indicate chance: the average number of fixations divided by the number of states (eleven). (E) Proportions of transitions (saccades) from the currently attended state (marked by an eye) to nearby states. The child proportions correspond to a single state (e.g. 36% of human transitions go to a child). (F) Illustration of frontier fixation/query. The red arrow indicates the next attended frontier state, and the red rectangle marks the path leading to the frontier state. (G) Probability of attending to a frontier state as a function of its path value (the sum of rewards on the path to that state). (H–I) Change in decision logit (H) and Q value (I) corresponding to a queried state and its ancestors, grouped by the queried state’s reward. Logits are normalized within sibling pairs to reflect the relative preference for a state over its sibling. Logits and Q values are baseline-corrected by subtracting the value immediately before the query (*t* = −1). (J–K) Change in the parent’s decision logit (J) and Q value (K) grouped by the revealed reward and the sibling’s value. K shows predictions from Bellman backups (Methods). Error bars show standard errors across participants or random seeds (and are often too small to be visible).

We trained the meta-RL agent on this task. Here, the mental actions correspond to querying a world model for the reward and local connectivity of a target state. Concretely, the information generator takes a state as input and returns (i) the associated reward, (ii) the *child* states that one could travel to from the target state, and (iii) the unique *parent* state from which one would reach the target. This mimics the visual information participants received with each fixation (the point value and arrows). Note, however, that similar architectures have been applied in cases where the world model resides in memory rather than a graphical display (e.g., Dyna; Sutton, 1990). To stabilize training and simplify analysis, we introduced two additional constraints. First, we required the network to plan and act in separate phases, introducing an additional action that switches from planning to action. Second, we prevented the network from querying a state before querying its parent. This forces the agent to plan forward, as is assumed by many algorithmic models of human planning (van Opheusden et al., 2023; Huys et al., 2015).

We then asked whether the agent displayed two key signatures of human behavior in the task identified by Callaway et al. (2024b): the tendency to fixate states close to (i) the initial state and (ii) the previously attended state. First, as illustrated in Figure 4D, both humans and the agent preferentially fixated/queried shallower states, as expected under non-exhaustive forward search. Notably, depth-1 states were attended more than once per trial; this is inconsistent with classical search algorithms (e.g., best-first search; van Opheusden et al., 2023) and optimal symbolic models (e.g., Sezener et al., 2019; Callaway et al., 2022) that perfectly represent a decision tree and search frontier. To test the second signature, we categorized pairs of consecutive fixations/queries based on the graphical relationship between the two states, specifically whether the next state was a child, parent, or sibling of the previous state (Figure 4E). Both humans and the agent showed a strong tendency to fixate/query children and siblings, as expected under breadth- or best-first search. Only humans, however, showed above-chance levels of parent fixations, potentially indicating explicit backtracking that the agent was more often able to avoid. Overall, these patterns suggest that both humans and the agent used strategies resembling tree search with a limited or noisy search frontier.

If both humans and the agent perform tree search, what guides their search? One likely candidate is “path value”: the cumulative reward along the path leading to a state (Figure 4F). Maximizing path value—or equivalently, minimizing path cost—is the default strategy in classical AI search (Dijkstra’s algorithm; Dijkstra 1959) and closely approximates rational metareasoning when a heuristic estimate of future value is unavailable, as in the current task. To test this hypothesis, we examined fixations to frontier states—states that had not yet been queried but whose parents had. Both humans and the agent preferentially attended to frontier states with higher path values (humans: *β* = .004, *p <* .001; agents: *β* = .009, *p <* .001; Figure 4G), and this tendency increased with higher search cost in the agent (Figure S3A-B). This suggests that both humans and the agent prioritize exploring promising paths to make efficient use of their limited resources. We further found that re-fixations/re-queries were directed toward states with higher Q values (i.e., expected cumulative future rewards) in both humans and the agent (humans: *β* = .006, *p <* .001; agents: *β* = .001, *p <* .001; Figure S3C), indicating a tendency to revisit states that had been discovered to yield good outcomes. This pattern intensified under memory constraints in the agent (Figure S3D), consistent with increased reliance on re-sampling to recover critical information. Together, these findings indicate that human planning reflects strategic computation selection shaped jointly by efficiency demands and memory limitations.

We then examined how each mental action updated the agent’s policy by analyzing the logits of decision actions during the planning phase (Figure 4H). Querying a state shifted its logit up or down depending on the sign and scale of the reward. The logit for the *parent* of the queried state shifted to a similar degree—but only if the queried reward was positive. These local updates closely tracked corresponding changes in Q values (Figure 4I), indicating that the agent learned to perform local value backups. Logits for further ancestors (e.g., the “grandparent;” middle panels) changed on the next timestep and to a weaker degree—notably, in a distance-independent manner. This suggests that the agent also tracked the path to each state (a “predecessor representation”; Bailey and Mattar, 2022). We further found that updates to the parent were gated by the sibling’s value: the parent’s logit increased primarily when the revealed reward exceeded the sibling’s value (Figure 4J). This is consistent with the maximization in the Bellman operator, where a state’s Q value depends only on the value of its best child (Figure 4K). However, in the cases when the revealed reward and sibling value were both low, the parent’s logit still did not decrease, in contrast to a full Bellman backup (see Figure S4 for a detailed comparison). This suggests that the learned backups were optimistic, propagating only positive updates up to the parent (surprisingly, this has little effect on behavior; Figure S5). Taken together, these results suggest that the agent learned an RL-like strategy for integrating feedback from queries, resonating with a previous proposal that planning can be viewed as stringing learning operations to construct action policies (Mattar and Daw, 2018).

### 2.5 Meta-RL agent reproduces human neural dynamics in a planning task

Lastly, we tested whether our framework could capture neural dynamics underlying human planning. A recent study by Vikbladh et al. (2024) provides an ideal testbed. Participants learned a nine-state graph arranged in a unidirectional loop. Each state had an associated object, which was defined by two features: color (yellow, green, or red) and category (vehicle, animal, or fruit) (Figure 5A). On each trial, participants received an offer specified by a start object (e.g., taxi), a reward feature (e.g., animal), and a stop feature (e.g., red) (Figure 5B-C). The total value of the offer was computed by traversing the graph from the start object until reaching an object with the stop feature, adding two points for each object matching the reward feature and subtracting one point for each non-matching object (Figure 5B). Participants performed the task in a magnetoencephalography (MEG) scanner. They completed a training phase and an immediate revaluation phase on the first day, followed by a second revaluation phase one week later (Methods).

**Figure 5:**
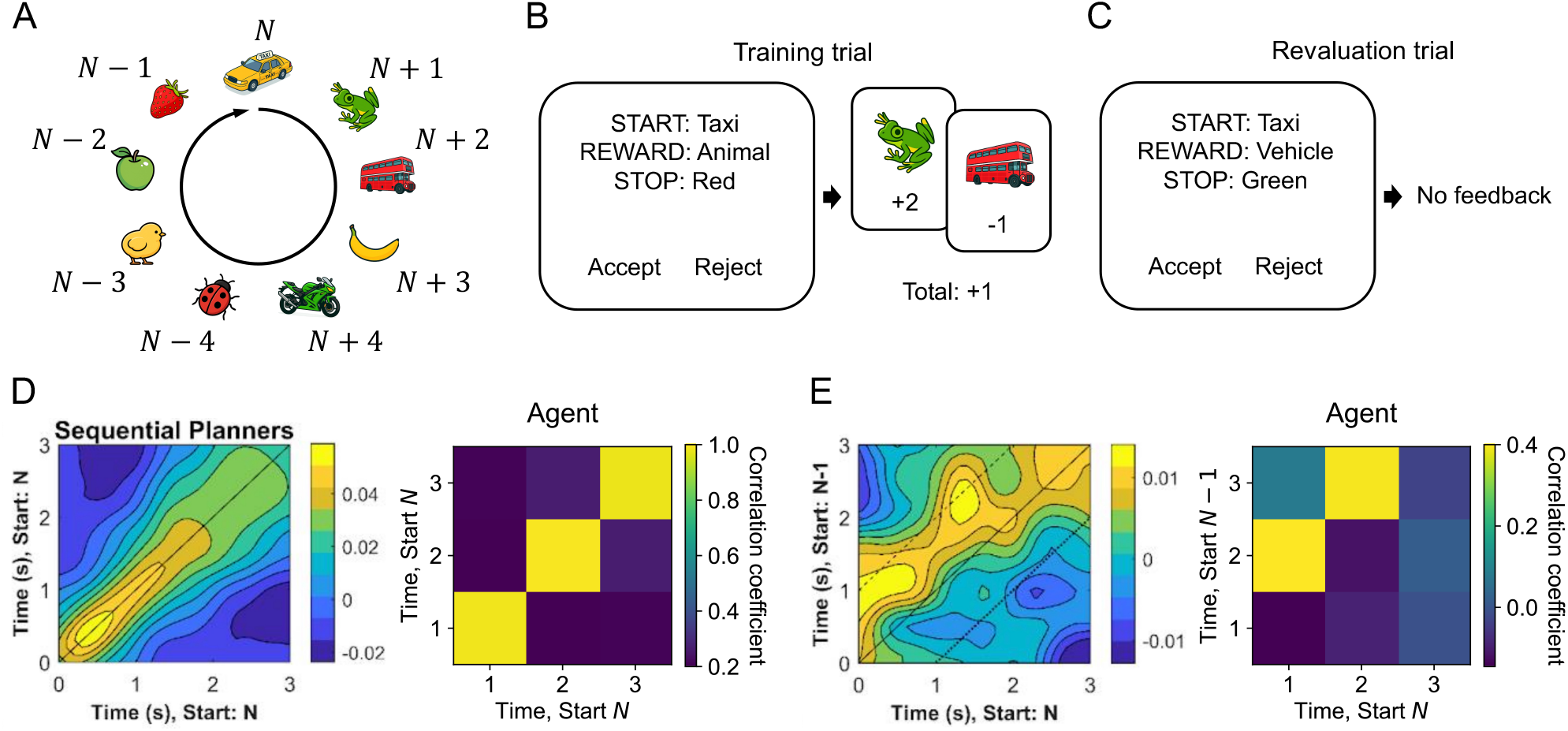
Meta-RL agent reproduces human neural dynamics in a planning task. (A) Graph structure. Nine states are arranged in a unidirectional loop. Each state is an object defined by two features: color (yellow, green, or red) and category (vehicle, animal, or fruit). *N* marks the start object. (B) Illustration of a training trial. Participants receive an offer specified by a start object (e.g., taxi), a reward feature (e.g., animal), and a stop feature (e.g., red), and they could choose to accept or reject the offer. The total reward of the offer is defined by traversing the graph from the start object until reaching an object with the stop feature, with each object matching the reward feature gaining +2 points and each object inconsistent with it getting −1 point. In the example, the total reward is +1. (C) Illustration of a revaluation trial. No feedback is given in revaluation trials. (D) Pattern similarity across voxels in sequential planners (left) and across hidden units in the agent (right), comparing trials with the same start object across time between halves. (E) Pattern similarity between trials starting at object *N* and trials starting at object *N* − 1. Task illustrations and panels showing experimental results are adapted from Vikbladh et al. (2024).

The key finding in this study was that brain activity during deliberation reflected step-by-step mental simulation. The authors identified participants whose choices are best explained by a model-based strategy and investigated whether they solved the task using “rollouts”, i.e., sequentially simulating future states along the graph. To test this, they split trials with the same start object (*N*) in half and computed correlations of activity across time points between halves. Correlations peaked along the diagonal, indicating stable dynamics for sequences with the same start object (Figure 5D, left). To examine if these dynamics reflected rollouts, the authors correlated trials with start object *N* against those starting one state earlier (*N* − 1). The authors found that the correlation was preserved but shifted up by a constant time lag, consistent with the hypothesis that neural dynamics reflected rollouts (Figure 5E, left). Notably, these dynamics were initially localized to the hippocampus but, one week later, were also found in the PFC, suggesting that planning-related representations are initially supported by the hippocampus and later consolidated into PFC.

We trained the agent on this task. We first assumed that a world model had already learned the task structure, which we did not model explicitly. Learning in the world model corresponds to the initial acquisition of task representations in the hippocampus. After the initial phase, the world model served as the information generator that produced synthetic experience to support consolidation (Girardeau et al., 2009; Girardeau and Zugaro, 2011; Ólafsdóttir et al., 2018). We then trained the agent on the synthetic data generated by the world model. This process parallels the slower consolidation occurring after the first day of the human experiment. The agent’s mental actions correspond to simulating the world model to query a state. Upon queried, the information generator returns the features of the queried state, as well as whether that state would yield a reward or trigger a stop (Methods). This setup enabled the agent to sequentially query the world model to perform mental simulations and decide whether to accept an offer based on simulated outcomes. We did not input the reward rule or stop rule directly to the agent; doing so would allow it to memorize the mapping from rules to decisions rather than solving the task through rollouts. This constraint reflects the view that PFC is not optimized to store task-specific input-output mappings, but instead learns abstract rules that generalize across tasks (Miller and Cohen, 2001; Wallis et al., 2001; Badre et al., 2010).

We asked whether the agent learned a rollout strategy. After training, the agent queried the next state along the graph from the current state 97% of the time, confirming that it had learned to perform rollouts. We then tested whether the agent exhibited the same rollout dynamics. We applied the same analysis to the agent’s hidden states (Methods). When correlating trials with the same start object, correlations peaked along the diagonal, indicating stable hidden dynamics (Figure 5D, right). When correlating trials offset by one start object, the peak shifted by one time step, mirroring the human MEG results (Figure 5E, right) and demonstrating that the agent’s recurrent dynamics reflect step-by-step mental simulation similarly in humans. Together, these findings provide a concrete example of how interactions between brain regions can be interpreted as implementing control of computations. Here, PFC learns to operate over hippocampal task representations to carry out computations that inform and refine future actions.

## 3 Discussion

In this work, we developed a neural network agent that learns to adaptively select computations. Adopting ideas in rational meta-reasoning, we formalized computations as mental actions: actions that leave the environment unchanged but generate decision-relevant information. We then trained an RNN with meta-reinforcement learning to integrate this information over multiple steps in order to effectively control both its external behavior and the computational process itself. The agent reproduced key signatures of the algorithms and representations used by both biological agents and optimal benchmarks. It selected computations consistent with human eye fixations, and it developed neural representations similar to those observed in primate PFC. Our work thus establishes a tractable and generalizable framework for modeling the adaptive control of computation and provides a theory of how such control can be implemented in neural systems.

Our key theoretical contribution is the union of two frameworks: meta-reasoning and meta-learning. The crucial bridge is the idea that reasoning can be understood as learning from information generated by one’s own cognitive operations (Matheson, 1968; Sutton, 1990; Lombrozo, 2024). With this view, each individual decision made by an agent is the result of an extended learning process. Learning how to reason is, in this sense, learning how to learn. Although this might strike some readers as a superficial terminological shift (or worse, a false equivalence; see “These points…” below), unifying these two frameworks requires non-trivial conceptual work on both sides—and it yields a framework more powerful and explanatory than each alone.

Classic meta-reasoning models assume a fixed computational architecture, allowing the agent to control which computations are executed, but not how they affect internal program state. Practically, this restriction makes it difficult to apply neural networks to meta-reasoning problems (Hay, 2016), limiting their application to simple, low-dimensional tasks. Theoretically, invoking a controller extrinsic to the core computational process raises concerns of infinite regress or relying on a “homunculus” to explain behavior without explaining the homunculus itself (Hazy et al., 2006; Botvinick and Cohen, 2014). Our framework addresses these challenges by redrawing the classical controller-architecture boundary. Specifically, we reduce the architecture to a stateless input-output module (the information generator) and combine integration and control into a single recurrent network. This allows a single system to jointly learn an algorithm and the representations it operates over.

Learning representations that support flexible context-sensitive learning and control is precisely the goal of meta-learning. However, these models focus on the interaction between an agent and its *external* environment, that is, learning about the world from observed data and controlling the world with overt behavior. Although some meta-learning models can be characterized as learning cognitive strategies for inference (Dasgupta et al., 2020) or decision-making (Binz et al., 2022), these strategies lack explicit structure and cannot adapt their computational effort based on the difficulty of individual problems. Recent work has begun to address this gap, but has been limited to binary arbitration between a simple and complex strategy (Moskovitz et al., 2024) or one-dimensional control over the amount of computation performed (Jensen et al., 2024). As a result, the computational strategies they can express are inherently limited in flexibility. Our framework addresses these limitations by re-conceptualizing the agent’s “environment” to include not only the outside world, but also the computational architecture of the agent itself (c.f. Simon, 1955). This allows the agent to adapt both the amount and nature of its computation to the structure of each problem it faces, in the same way that a standard meta-learner adapts its behavior. Because (mental) action sequences are inherently combinatorial, this form of computational adaptation can generalize systematically in a way that undifferentiated distributed systems have failed to.

These points notwithstanding, one could still reasonably ask whether modeling computation as active learning amounts to falsely equating two fundamentally different things. This concern is sharpened by the fact that some of our case studies used gaze as an index of supposedly internal computation. However, it is ameliorated by several factors. First, our framework also applied in settings where mental actions did not map to gaze (Figure 3 A-E) and where mental actions corresponded to objects that were not currently visible (Figure 5). Second, gaze is often interpreted as an indicator of (or anchor for) internal processing. This interpretation is easily justified in the simple choice tasks, where the stimuli were either familiar snacks (Figure 2) or heavily trained cues (Figure 3) whose identity could be perceptually resolved far faster than the observed response times (and without attentional alternation). Admittedly, in the planning task (Figure 4), external information gathering clearly plays a substantial role. However, the observed fixations showed clear signatures of a computational process (planning) that would be difficult to explain if they were not accompanied by internal operations (e.g., summing the rewards along a path; Figure 4G). Finally, although internally generated and externally perceived information may differ in important ways (e.g., accessibility and reliability), the fact that we were able to successfully model both with one abstraction—the information generator—suggests that their functional roles may be quite similar (c.f. Hunt et al., 2021). Recognizing this shared functional role may help us understand more naturalistic decision-making scenarios, where both internal and external information sources must be integrated and sought out in a tight, interdependent loop.

In our framework, we modeled a feedback loop between an RNN and an information generator. Biologically, we interpret this loop as mapping onto interactions between PFC and other brain regions. In line with this view, PFC has been shown to maintain and update belief states (Chan et al., 2016; Schiereck et al., 2025; Wilson et al., 2014; Schuck et al., 2016; Bartolo and Averbeck, 2020; Starkweather et al., 2018), whereas regions such as the hippocampus and basal ganglia are well positioned to serve as information generators that perform task-specific computations and feed information back to update representations in PFC (Jensen et al., 2024; O’Reilly and Frank, 2006; Wassum, 2022; Stopper et al., 2014; Pemberton et al., 2024). Recent work has begun to couple RNNs with world models or memory systems to model how prefrontal–hippocampal interactions implement planning and memory retrieval (Jensen et al., 2024; Lu et al., 2022; Li et al., 2024). Our work contributes to this growing line of research by providing a general framework for specifying such models and revealing their deep connections to normative theories of cognition.

We modeled the information generator as a fixed module that implements a predefined set of computations, leaving open how the effects of computations are acquired. One natural extension is to explicitly model the information generator and jointly optimize both the controller and the effects of computations. With this approach, computations themselves could adapt to task structure and the controller’s demands, enabling the study of how computations are acquired, refined, and composed across contexts (e.g., Chang et al. 2018). For example, in hierarchical planning, computations operate at different levels of abstraction, and humans flexibly plan at the appropriate granularity (Correa et al., 2023; Tomov et al., 2020; Solway et al., 2014). Jointly optimizing the information generator could allow the agent to learn computations at multiple levels of abstraction.

Although our focus has been on capturing the flexibility and efficiency of neural processing in humans and other primates, our work also speaks to basic questions about intelligence broadly construed. Extreme flexibility has long been touted as a unique feature of human intelligence, but recent advances in AI have put this view on increasingly shaky ground. Crucially, however, these advances have been coupled with a massive increase in computation—when it comes to efficiency, the brain remains the clear leader. By showing how flexible and efficient computation can be implemented in a brain-like recurrent architecture, we hope to have provided some hints about how these two desirable features could be simultaneously reflected in artificial systems.

## 4 Methods

### 4.1 Meta-RL agent

#### 4.1.1 Network architecture

The meta-RL agent was implemented as a fully connected gated recurrent unit (GRU) network with an actor-critic architecture. The hidden state of the GRU was fed into two linear output heads: the policy head and the value head. The policy head took the hidden state *h*_*t*_ as input and produced action logits, which were passed through a softmax function to produce the policy *π*_*θ*_(*a*_*t*_ | *h*_*t*_) from which actions were sampled. The value head took the hidden state *h*_*t*_ as input and produced the value baseline *V*_*θ*_(*h*_*t*_).

#### 4.1.2 Model training and evaluation

The agent was trained with REINFORCE with a baseline to maximize the expected cumulative reward. The total loss ℒ was a weighted sum of the policy loss ℒ_*π*_, the value loss ℒ_*v*_, and the entropy regularization loss ℒ_*e*_:

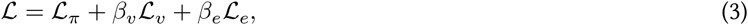

where *β*_*v*_ and *β*_*e*_ are the weights and the individual losses are defined as:

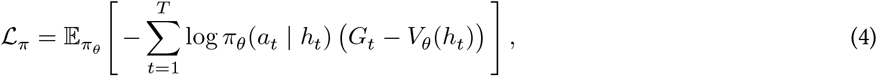

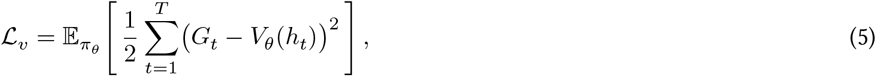

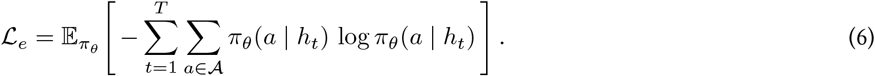

Here, *a*_*t*_ is the action selected at time *t*, 𝒜 denotes the set of valid actions, and *G*_*t*_ is the empirical return from time step *t*, that is, the sum of rewards from that point onward: 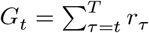

Unless otherwise noted, the agent’s hidden states were set to zeros at the beginning of an episode. The agent was trained with the ADAM optimizer (Kingma and Ba, 2014) using a batch size of 40 and a learning rate of 10^−3^, with episodes independently sampled from all possible environment configurations. We set the weights of the value and entropy losses (*β*_*v*_ and *β*_*e*_) separately for each case study to ensure training stability and roughly match the degree of noise in the behavior/neural recordings (the final value of *β*_*e*_ is akin to a softmax temperature). To further enhance training stability, gradients were clipped to a maximum norm of 1.0.

During evaluation, model parameters were fixed and no further learning took place. Instead, adaptations were achieved through the agent’s hidden dynamics. Actions were sampled on-policy. All results were aggregated over 5 agents trained with different random seeds, with each agent simulating at least 10^5^ episodes.

#### 4.1.3 Action masking

In some simulations, we applied action masking to ensure that only a subset of valid actions was allowed to be executed in specific time steps. Concretely, we set the logits of masked actions to negative infinity. This ensured that their probabilities under the policy were 0 so they would never be sampled. Additionally, when computing the entropy loss, their corresponding term *π*_*θ*_(*a* | *h*_*t*_) log *π*_*θ*_(*a* | *h*_*t*_) was manually set to 0. As a result, the agent could not adjust the probabilities of masked actions to influence the entropy loss.

### 4.2 Simple choice task in Callaway et al. (2021)

#### 4.2.1 Task setting and model training

In the task, the agent was presented with 2 or 3 items whose true values were unknown, and had to gather noisy value samples through sampling actions and select the item with the highest value. The agent was based on a GRU network with 48 units. The input consisted of the identity of the currently attended item, the observed value sample from that item, the previous action, and the previous reward. The identity of the attended item and the previous action were encoded as one-hot vectors. The action space consisted of either 4 or 6 actions, corresponding to 2 or 3 sampling actions and 2 or 3 decision actions, for binary and trinary choice task, respectively. At the beginning of each episode, the attention pointer was initialized to blank (corresponding to all zeros in the one-hot encoding) and the value sample was set to 0. Executing a sampling or decision action moved the pointer to the corresponding item. Episodes terminated when the agent executed a decision action to choose an item or when the maximum time horizon of 100 steps was reached.

In the original experiments (Krajbich et al., 2010; Krajbich and Rangel, 2011), participants rated 70 snack items on a −10 to 10 scale, and these ratings served as the true values of the items. Participants then completed 100 choice trials with only items rated non-negatively. Previous work has suggested that the initial rating phase (which includes both positively- and negatively-rated items) may influence participants’ prior beliefs about item values (Callaway et al., 2021; Frömer et al., 2025). To instill this prior into the meta-RL agent, we wanted to train the agent on the same value distribution. However, since the experiments only included items with non-negative ratings and those with negative ratings were unavailable, we did not have access to the full prior distribution. Instead, we estimated the distribution by fitting a truncated Gaussian (restricted to non-negative values) to the empirical value distribution from the binary choice task (Krajbich et al., 2010) using maximum likelihood estimation. We then trained the agent using the full Gaussian distribution (including both non-negative and negative values) with the same parameters as the truncated fit. This distribution had a mean of 0.087 and a standard deviation of 4.33. In each episode, the true values were randomly sampled from this Gaussian distribution and rounded to the nearest integer.

At each sampling step, the value sample from the currently attended item was drawn from a Gaussian distribution centered on the item’s true value, with a standard deviation of 7. Each sampling step incurred a cost: 0.005 for maintaining attention on the same item and 0.12 for switching to a different item (c.f. Callaway et al., 2021). The agent was trained for 1.2 × 10^7^ episodes. For training stability, all rewards were scaled by a factor of 0.2, which did not affect the optimal policy. We set the value loss coefficient *β*_*v*_ to 0.05, and linearly annealed the entropy loss coefficient *β*_*e*_ from 0.05 to 0.015 in the binary choice task and from 0.05 to 0.01 in the trinary choice task.

#### 4.2.2 Model evaluation

When evaluating the agent’s behavior (Figure 2C-F) and decoding belief states from its hidden states (Figure 2G), we simulated the agent using only items with non-negative ratings, consistent with the original experiments. When performing PCA on the hidden state (Figure 2K–N), we simulated the agent with both non-negative and negative ratings, which better covered the agent’s experience during training. For PCA visualization, we simulated the model with the standard deviation of the value-sampling distribution set to 0, so that samples were identical to the true values. This allows one to see the grid structure and order invariance inherent in the space, uncorrupted by noise in realized value samples.

#### 4.2.3 Belief state decoding

We trained linear decoders to decode sufficient statistics of normative belief states from the agent’s hidden states (Figure 2G). Posterior means and precisions were computed by conjugate Gaussian inference with known observation noise, following Callaway et al. (2021). We initialized the prior distribution as a Gaussian with the same parameters used to train the agent. At each time step, the agent’s value sample was incorporated via standard Bayesian inference to update the belief of the corresponding item. This process yielded the belief state sequences predicted by the optimal symbolic model. We then trained linear regression decoders to decode the Bayesian means and precisions as well as the attended item from the hidden states. Decoding performance was evaluated using 5-fold cross-validation, averaged across 5 random seeds and across 2 or 3 items.

### 4.3 Simple choice task in Rich and Wallis (2016)

#### 4.3.1 Task setting and model training

The model was similar to the binary choice task described earlier with some modifications to capture the different task structure. Most notably, we masked decision actions for the first 15 time steps. This roughly captures the fact that macaques were required to maintain fixation on an option for at least 450 ms before committing a choice. True item values were sampled uniformly from the set {1, 2, 3, 4}. We set the standard deviation of the Gaussian sampling distribution to 1 and the costs to 0.07 and 0.1 for sampling the same/different item, respectively (the difference in cost structure captures the faster response times and more rapid alternation). As before, the agent was trained for 1.2 × 10^7^ episodes. All rewards were scaled by a factor of 0.5. We set the value loss coefficient *β*_*v*_ to 0.05, and linearly annealed the entropy loss coefficient *β*_*e*_ from 0.05 to 0.01.

#### 4.3.2 Linear discriminant analysis

In the original study, the authors trained a linear discriminant analysis (LDA) model to classify the value (i.e., learned reward association) of images presented in isolation, and then applied this model to decode value representations during binary choice trials. Because our model does not distinguish between an item’s identity and its value and we have direct access to the agent’s attention, we instead trained an LDA model directly on choice trials. Concretely, we trained the LDA model to classify the value of the currently attended option from the network’s hidden state immediately after receiving the value sample, using 5-fold cross-validation. In the experiments, neural activity was decoded using an 800 ms window following picture onset. Given that the median latency from picture onset to the choice-initiating fixation was 223–224 ms, the window thus spanned both pre-choice and post-choice periods. For the agent, however, post-choice time steps carry no meaningful information. We therefore restricted decoding to the interval between picture onset (episode start) and the time step at which the choice was made. Finally, in the experiment, a value state was defined when the posterior probability from the LDA decoder exceeded 0.5 for at least 35 ms. For the agent, we adopted an analogous criterion, requiring the posterior to exceed 0.5 for at least one time step. A transition between value states was defined as a shift from the value state of one option to that of the other.

#### 4.3.3 Regression analyses

We used linear mixed-effects models to evaluate the relationship between the number of transitions from one option value to the other and the values of both options. We fit a mixed-effects model with random intercepts and slopes across sessions or random seeds:

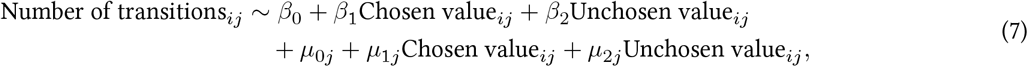

where *i* indexes trials and *j* indexes sessions or random seeds. Here, *β*_0_, *β*_1_ and *β*_2_ are the fixed intercept and slope, while *µ*_0*j*_, *µ*_1*j*_ and *µ*_2*j*_ represent session- or seed-specific random effects.

### 4.4 Simple choice task in McGinty and Lupkin (2023)

#### 4.4.1 Task setting and model training

The task was identical to the previous task except that true values were sampled from the set {1, 2, 3, 4, 5}, the standard deviation of the sampling distribution was set to 5, and the costs were set to 0.008 and 0.25 for sampling the same/different option, respectively. All rewards were scaled by a factor of 0.2. The network structure and training procedure were unchanged.

#### 4.4.2 Model evaluation

For this experiment, we were specifically interested in characterizing the network’s learned representational dynamics. To isolate those dynamics, we simulated the network with a fixed attentional policy. Concretely, we required the agent to sample one item for two steps and then the other item for two time steps, with the order randomized across trials. We recorded the initial hidden state and the four subsequent states. This procedure isolates representational effects from sampling policy. It ensures equal coverage of all value pairs in the simulated dataset to prevent sample durations from being influenced by option values (as observed in Figure S1G–J), and guarantees temporal alignment across trials.

#### 4.4.3 Value subspace analysis

We applied the value subspace analysis of McGinty and Lupkin (2023) to the macaque OFC neural recordings and the agent’s hidden states (Figure 3F-G). For macaques, we used neural activity from 200 ms before to 500 ms after the second fixation, using 200 ms bins and 100 ms increments. For the agent, we used the hidden activity of the first 5 time steps. We concatenated the activity from each time bin or time step into a data matrix 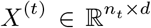, where *n*_*t*_ is the total number of trials within a session or a seed, and *d* is the dimensionality of the activity. For each session or seed, the data were first split into a training set and a test set. We identified value subspaces using only the training set, and performed subsequent analyses on the test set. To identify value subspaces, we trained LASSO decoders to predict the values of the first- and second-viewed options from either OFC activity or the agent’s hidden states during the period when both value representations were relatively stable, specifically, the mean activity from 200 to 400 ms after the second fixation for the macaque, and the final time step for the agent (denoted 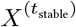). The regularization parameter of the LASSO decoders was optimized within the range 0.001 to 10 to minimize the 10-fold cross-validation error on the training set. The resulting decoder weights for the first- and second-viewed options (*β*_1_ and *β*_2_) defined a two-dimensional value subspace residing in the *d*-dimensional neuron/unit space.

To analyze and visualize dynamics within the value subspace, we transformed the decoder weights into an orthonormal basis for the value space, using the same method as McGinty and Lupkin (attributed to Semedo et al. (2019)). This method identifies a *d ×* 2 matrix *Q* whose columns are orthogonal, lie in the span of [*β*_1_ *β*_2_], and are most closely aligned with the covariance structure of the dataset 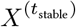. Concretely, we defined *B* = [*β*_1_ *β*_2_] and 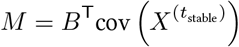. We then performed a singular value decomposition, *M* = *UDV* ^T^ and define *Q* as the first two columns of *V*. See Semedo et al. (2019) for further explanation of the method.

We then projected neural activity from each time step or time bin onto the identified subspace, 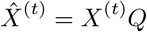. The value subspace analysis was conducted separately for each session or random seed. In the macaque data, 4 sessions were excluded because the decoders failed to reliably decode either value (i.e., cases where weights collapsed to all zeros for at least one decoder), leaving a total of 28 sessions for analysis.

To account for the temporal lag in neural processing not captured by our model, we aligned the two time courses differently. For macaques, we aligned neural activity to the start of the second fixation, which occurred about 200 ms after the first fixation. Under this alignment, value signals associated with the first-viewed option began to emerge around *t* = 0 ms, while signals for the second-viewed option began to emerge roughly 200 ms later (*t* = 200 ms). For the agent, sampling effects influenced the hidden state immediately, so trials were instead aligned to the start of the trial. With this alignment, the influence of sampling the first item emerged after the first time step (*t* = 1), and the influence of sampling the second item emerged after the third time step (*t* = 3). This process ensured that temporal dynamics were comparable between the macaque and the agent.

#### 4.4.4 Decoding values from the value subspace

To quantify the temporal emergence of value representations within the value subspace, we performed decoding on the projected data separately at each time bin or time step using the same LASSO decoding procedure. This yielded a time-resolved measure of value-decoding performance within the value subspace (Figure 3I). We evaluated decoding using the projected rather than the raw data because our goal was to characterize how the value gradient unfolded specifically within the value subspace over time.

#### 4.4.5 Rotation analysis

To identify rotation angles (Figure 3J-K), we first repeated the decoding analysis separately at each time bin (for the macaque, starting from *t* = 0 ms) or time step (for the agent, starting from *t* = 1). This yielded two weight vectors per time bin or time step, 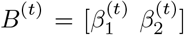. We then wanted to find the direction of the value gradient in neural activity for each time bin or time step. Note that the raw decoder weights *B*^(*t*)^ do not directly correspond to the direction along which neural activity changes as a function of value. Instead, the direction of the value gradients in neural activity is given by the covariance-weighted (encoding) directions *B*^(*t*)T^cov (*X*^(*t*)^). Accordingly, we transformed the decoding weights into encoding directions by multiplying them with the covariance matrix, *M*^(*t*)^ = *B*^(*t*)T^cov (*X*^(*t*)^). We then projected these encoding directions onto the previously identified orthogonal basis of the value subspace (at *t*_stable_), 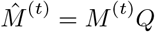. The resulting projected row vectors in *M* ^(*t*)^ characterize, at each time bin or time step, the direction of the value gradient for the first-viewed and second-viewed option within the value subspace. Rotation angles were then obtained by computing the angle of each row vector and subtracting the first-viewed option’s gradient angle at *t* = 0 ms for macaques and *t* = 1 for the agent (Figure 3K).

### 4.5 Planning task in Callaway et al. (2024b)

#### 4.5.1 Task setting and model training

In the task, human participants were presented with a decision tree displayed as a graph (Figure 5A). They could freely view the tree to collect information, and then chose a path by sequentially clicking states on the path, aiming to maximize rewards. The decision tree was randomly sampled from all possible binary trees with 11 states, and rewards were uniformly sampled from {−8, −4, −2, −1, 1, 2, 4, 8}. All the rewards and transition structures are randomized every trial, ensuring decision-time planning. Each trial had a 15-second time limit, indicated by a collapsing bar beside the tree.

The agent was implemented as a GRU network with 128 units. Its input consisted of the identity of the currently attended state, its parent and two children, the identity of the physical state, the reward of the attended state, the trial phase, elapsed time, the previous action, and the previous reward. All the state identities as well as the previous action were encoded as one-hot vectors, yielding a total input dimensionality of 82. The action space contained 11 query actions, 11 decision actions, and one phase-transition action, yielding 23 actions in total. Each query or decision action corresponded to a specific state. Executing the query action moved the attention pointer to that state, while executing a decision action moved the both the physical state and the attention pointer to that state. At the start of an episode, the attention pointer was initialized to the physical state (the root).

Each episode consisted of two phases: a planning phase and a decision phase. During the planning phase, only query actions and the phase-transition action were available, while decision actions were masked. Moreover, we applied action masking to make sure that a state could be queried only after its parent had been queried. This constrained the agent to forward search, as is assumed by most models of human search (van Opheusden et al., 2023; Huys et al., 2015) and strongly reflected in the gaze data. Importantly, this constraint did not prevent re-queries (e.g., returning to a parent) or jumps within the tree (e.g., to a sibling). In the decision phase, only decision actions were allowed. At each time step, the agent could only select one of the two children of the current physical state, consistent with the experimental task. Episodes terminated when the agent reached a terminal state or when the maximum horizon of 100 steps was exceeded.

Each query action incurred a cost of 0.24. The agent was trained for 1.5 × 10^7^ episodes. All rewards and costs scaled by a factor of 1*/*8. The value loss coefficient *β*_*v*_ was set to 0.05, and the entropy loss coefficient *β*_*e*_ was linearly annealed from 0.05 to 0.04.

#### 4.5.2 Memory manipulation

We injected noise into the agent’s hidden state to manipulate its memory capacity (Figure S3). To ensure that the variance of the hidden state was preserved, we added noise through:

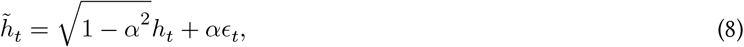

where *h*_*t*_ and 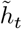 denote the hidden state before and after noise injection, *ϵ*_*t*_ is the random noise drawn from a Gaussian distribution with mean 0 and variance matched to that of *h*_*t*_, and *α* controls the noise proportion. With this setting, a fraction of the variance in the hidden state was replaced with random noise at each time step. Intuitively, larger values of *α* forced the agent to forget more information.

#### 4.5.3 Fixation/Query data analyses

For all analyses of human fixation data, we only included fixations prior to the first click, and further excluded fixations immediately preceding the first click, as these fixations likely guided motor execution rather than reflecting planning. For the agent, we included all querying steps before the agent executed the phase-transition action.

#### 4.5.4 Regression analyses

We used linear mixed-effects models to evaluate the relationship between the probability of fixating/querying frontier states and their path values (Figure 4G, Figure S3A) and the relationship between the probability of re-fixating/re-querying a state and its Q value (Figure S3C). For frontier fixations/queries, we first filtered all queries directed to frontier states. For each such query, we considered every possible frontier state, recorded its path value, and coded whether it was fixated/queried (1 if fixated/queried, 0 otherwise). We then fit a mixed-effects model with random intercepts and slopes across participants (or random seeds for the agent):

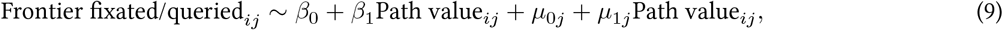

where *i* indexes frontier states and *j* indexes participants or random seeds. Here, *β*_0_ and *β*_1_ are the fixed intercept and slope, while *µ*0*j* and *µ*_1*j*_ represent participant- or seed-specific random effects.

For re-fixations/re-queries, we computed the Q value for every state that had been fixated/queried before. We coded whether each of these states was fixated/queried at the current time step (1 if fixated/queried, 0 otherwise), and fit the following mixed-effects model with random intercepts and slopes across participants (or random seeds for the agent):

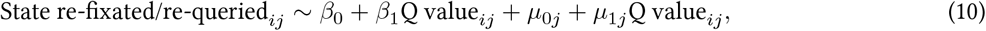

where *i* indexes possible children in each event and *j* indexes participants or random seeds.

When evaluating how these two relationships varied with cost or noise, we instead fit linear regression models to all episodes from each agent trained with a given random seed and then averaged the resulting slopes across seeds (Figure S3BD).

#### 4.5.5 Backup analyses

We computed ground-truth Q values at each query step given the explored tree using dynamic programming, with unqueried states assigned a reward of 0. We then compared query-induced changes in Q values and compared them against the corresponding changes in the agent’s decision logits (Figure 4H-I).

To investigate the agent’s learned backup strategies, we identified steps in which a state was queried for the first time while its sibling had already been queried, and measured the resulting change in the decision logit of their parent. We benchmarked these updates against Bellman backups by simulating how Q values would change under the Bellman equation (Figure 4J-K; Figure S4). For a queried state *s* with parent *s*^parent^ and sibling *s*^sibling^, the parent’s Q value was updated as:

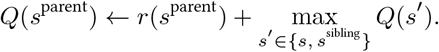

For the optimistic Bellman update, we additionally clipped the updated value at zero:

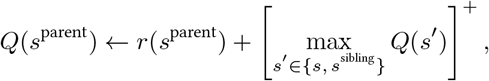

where *x*^+^ ≡ max(*x*, 0).

### 4.6 Planning task in Vikbladh et al. (2024)

#### 4.6.1 Task setting and model training

In the task, participants learned a nine-state graph arranged in a unidirectional circle, where each state was an object defined by a color and a category. For each participant, one feature dimension indicated reward and the other indicated when to stop. In each trial, participants made an accept-reject decision for a presented offer. The total reward of the offer was computed by traversing the graph from the start object until reaching an object with the stop feature. If the cumulative reward along the sequence (rewarded objects +2, non-rewarded −1) is positive, they should accept; otherwise, they should reject. The experiment included a training phase followed by immediate and delayed (seven-day) revaluation phases. During training, participants completed 20 repetitions for each start object, made accept/reject decisions, and received feedback showing the traversed sequence and resulting points. The reward feature and stop feature were fixed per start object. In each revaluation phase, every start object was paired with all combinations of reward and stop features, yielding 81 offers (72 novel), and no feedback was provided.

The agent was implemented as a GRU network with 64 units. It maintained an attention pointer, which was initialized to the start state. Its input consisted of the identity of the start state, the identity of the currently attended state, the reward feature and stop features of the attended state (e.g., “yellow” or “vehicle”), binary indicators for whether the currently attended state was rewarded or marked a stop, and the previous action and reward. State identities and features were one-hot coded. The action space comprised 11 actions: 9 simulation actions (one per state), an accept action, and a reject action. When the agent selected a simulation action, it queried the world model, which updated the attention pointer to the queried state, generated a prediction of that state’s features, and returned them to the agent on the following time step. A trial ended when the agent executed accept or reject actions.

To prevent the agent from learning a stereotyped strategy, we trained the agent on all possible combinations of reward and stop features instead of only using fixed reward and stop features per start object. Each simulation action incurred a cost of 0.02. The agent was trained for 1 × 10^7^ episodes. Because the agent has no prior knowledge of which states lie on the path from the start to the stop state, credit assignment can be challenging. To address this, we applied reward shaping early in training to discourage simulation of off-path states. Concretely, each simulation step on an off-path state incurred a penalty linearly annealed from 0.1. to 0 over the first 25% of training. All rewards were scaled by a factor of 0.5. The value loss coefficient *β*_*v*_ was set to 0.05, and the entropy loss coefficient *β*_*e*_ was linearly annealed from 0.05 to 0.005. With all-zero initialization, hidden states at the start of every episode would be identical, leaving the subsequent similarity analysis undefined at the first time step; we therefore initialized hidden states by sampling from a unit Gaussian.

#### 4.6.2 Similarity analysis

We conducted the same similarity analysis on the agent’s hidden dynamics (Figure 5D-E). In the original study, similarity was evaluated over the first 3 seconds; here, we examined the first three time steps of each trial. For each start object, we randomly split trials either with the same start object (*N* vs. *N*) or with a one-object lag (*N* vs. *N* −1) into two halves. We then Z-scored hidden states for each time step within each half and computed the correlation of hidden activity across time steps between the two halves. For human participants, this procedure was repeated for 50 random splits per start object per participant, and correlations were averaged across start objects and participants. For the agent, we repeated this procedure for 1000 random splits per start object per random seed.

### 4.7 Code and data availability

Code for this paper is available at https://github.com/SixingChen028/recurrent-meta-reasoning. Human behavioral data from Krajbich et al. (2010), Krajbich and Rangel (2011), and Callaway et al. (2024b) are included in the repository. Macaque electrophysiology data from Rich and Wallis (2016) and McGinty and Lupkin (2023) are available upon request to the original authors.

## 5 Acknowledgements

We thank Ian Krajbich for generously sharing data. We are grateful to Kristopher T. Jensen and the Mattar lab for valuable feedback on the project and manuscript. The original work of Rich and Wallis (2016) was supported by NIMH grants R01-MH097990 and R01-MH131624.

## 6 Supplementary Information

### 6.1 Supplementary figures for the simple choice task in Callaway et al. (2021)

**Figure S1:**
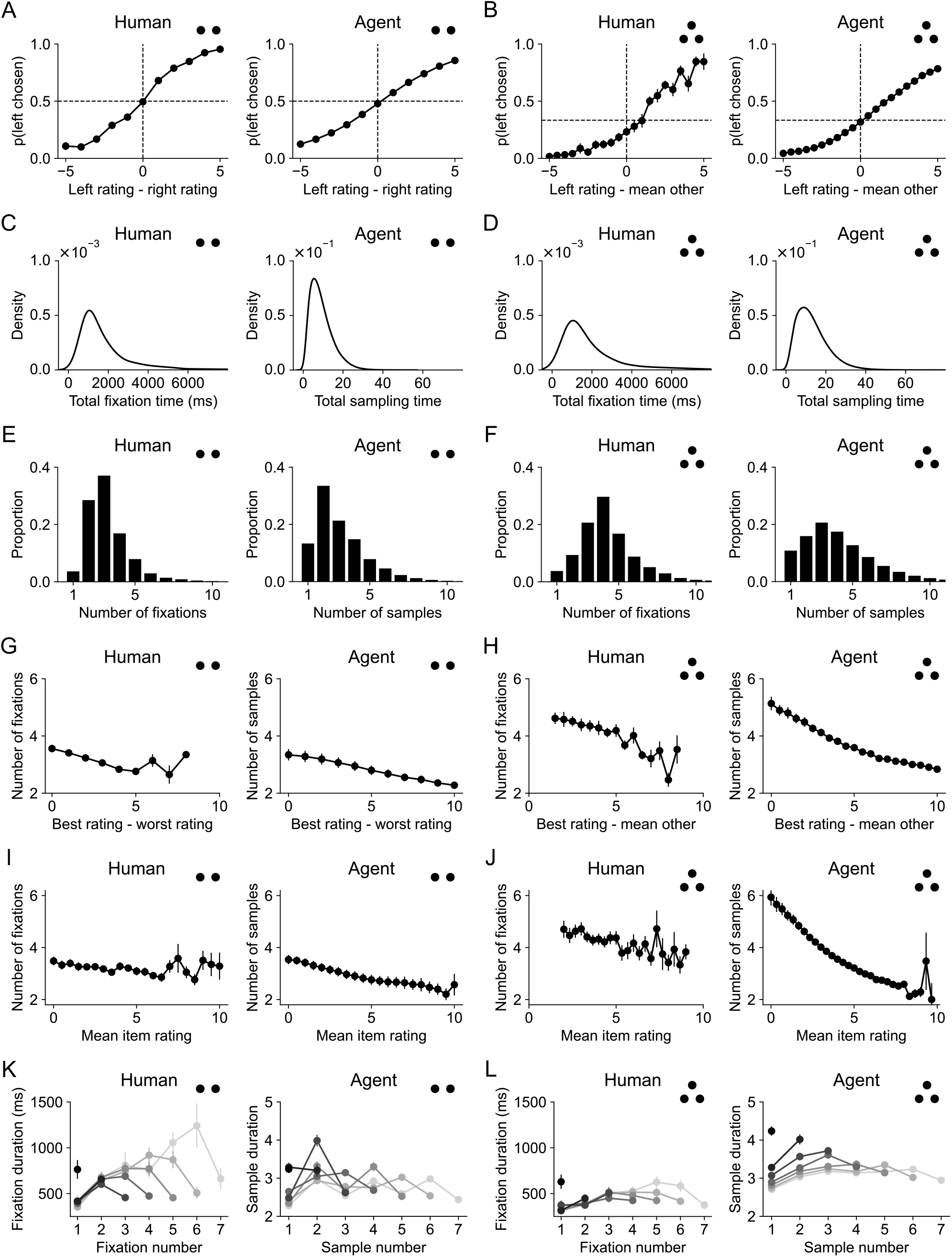
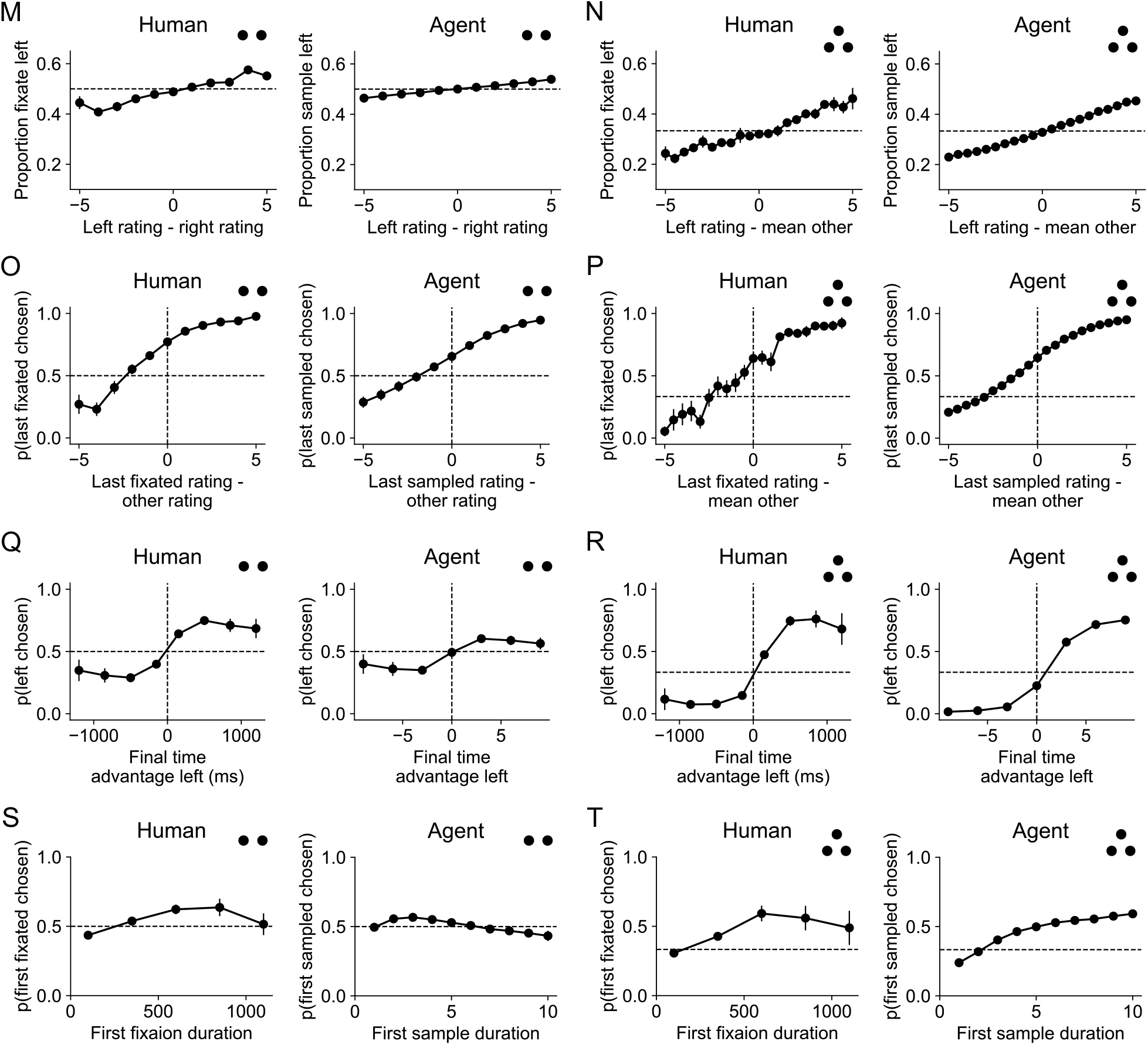
Behaviors from humans and the meta-RL agent on the simple choice task. (A–B) Choice probability as a function of relative rating. (C–D) Distribution of total fixation or sampling time. (E–F) Histogram of number of fixations or samples in a trial. Here a sample refers to a contiguous period of sampling from the same item, analogous to a fixation in humans. (G–H) Number of fixations or samples as a function of relative rating. (I–J) Number of fixations or samples as a function of mean rating. (K–L) Fixation or sample duration as a function of fixation or sample number. Colors correspond to number of total fixations or samples within a trial. (M–N) Proportion of time attending to the left item as a function of its relative rating. (O–P) Probability that the last attended item is chosen as a function of its relative rating. (Q–R) Probability that the left item is chosen as a function of its final time advantage, given by total fixation or sampling time to the left item minus the mean total fixation or sampling time to the other item(s). (S–T) Probability of choosing the first fixated item as a function of first fixation or sample duration.

### 6.2 Supplementary figures for the simple choice task in Callaway et al. (2021)

**Figure S2:**
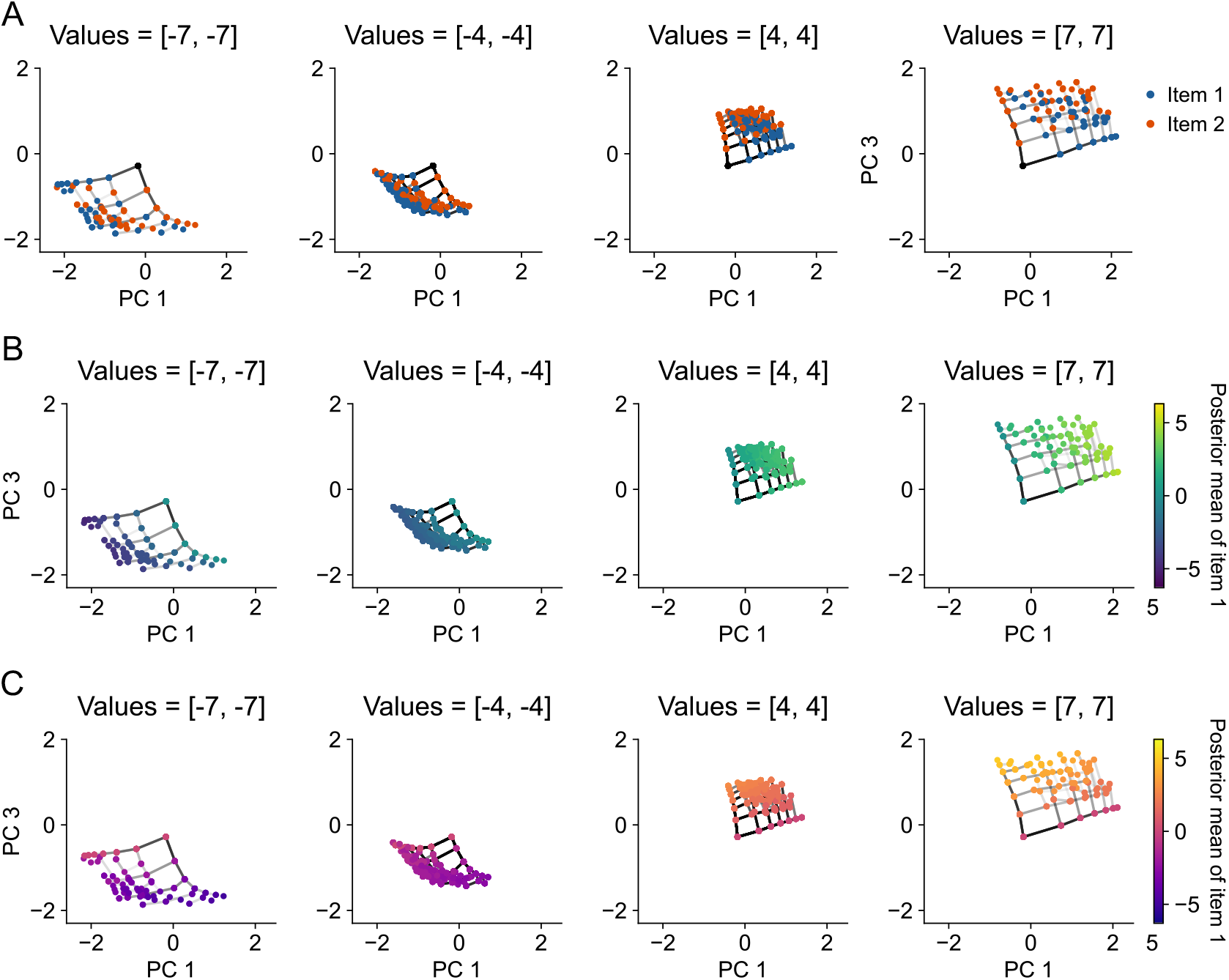
PCA results for different value conditions in the binary choice task. Hidden states are colored by the current attended item (A), posterior mean of item 1 (B), and posterior mean of item 2 (C). Panels show conditions where both items have values of -7, -4, 4, and 7, respectively. Line transparency shows transition frequency. Hidden states are simulated with noise-free value samples for visualization (Methods).

### 6.3 Supplementary figures for the planning task in Callaway et al. (2024b)

**Figure S3:**
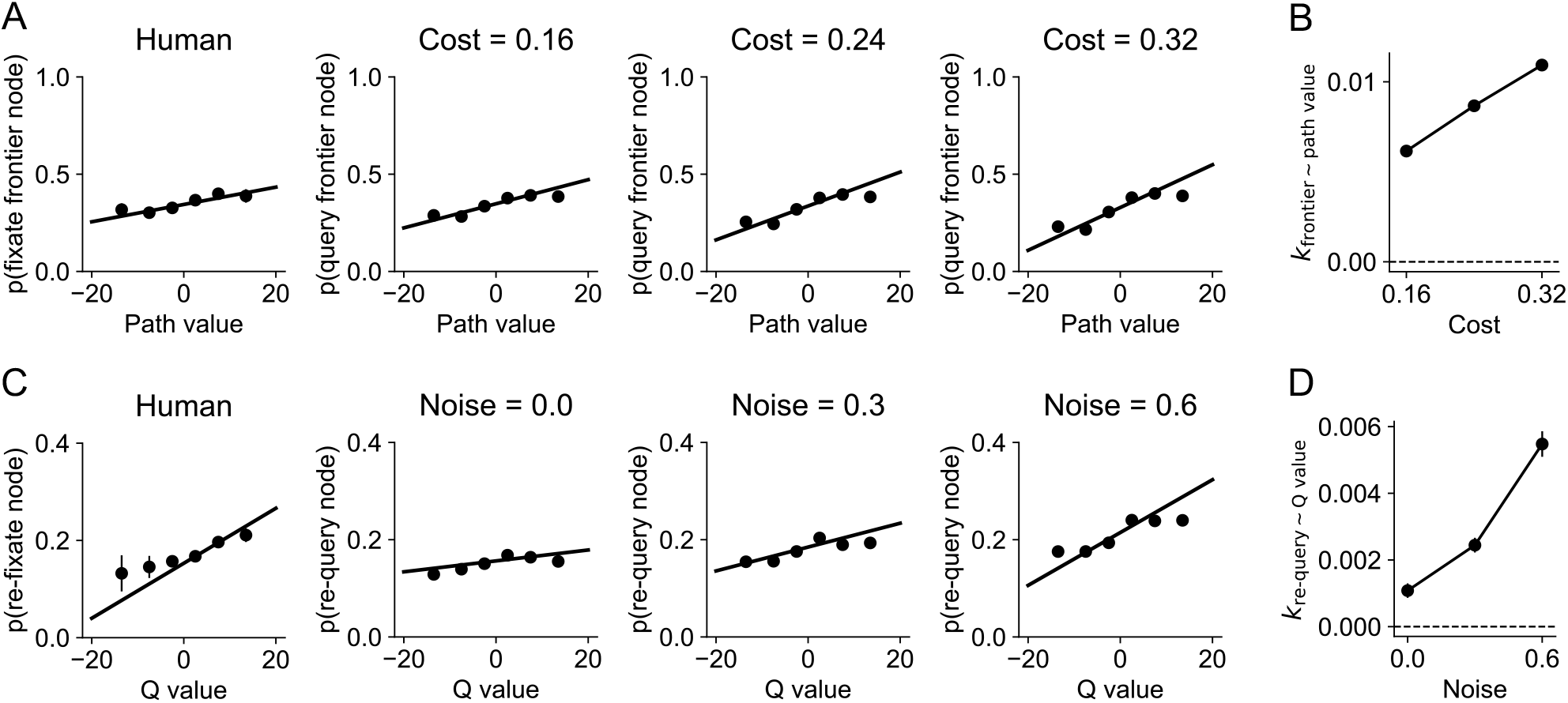
Planning strategy under different levels of search cost and memory capacity. (A) Probability of attending to a frontier state as a function of it path value for humans and agents with different costs. (C) Correlation coefficient between frontier fixation/query and path value as a function of cost per query step. (E) Probability of re-attending to a state as a function of its Q value for humans and agents with different noise levels in hidden states. (F) Correlation coefficient between re-fixation/re-query and Q value as a function of noise. Here noise refers to the parameter *α* that specifies the proportion of hidden state variance replaced by noise (Methods).

**Figure S4:**
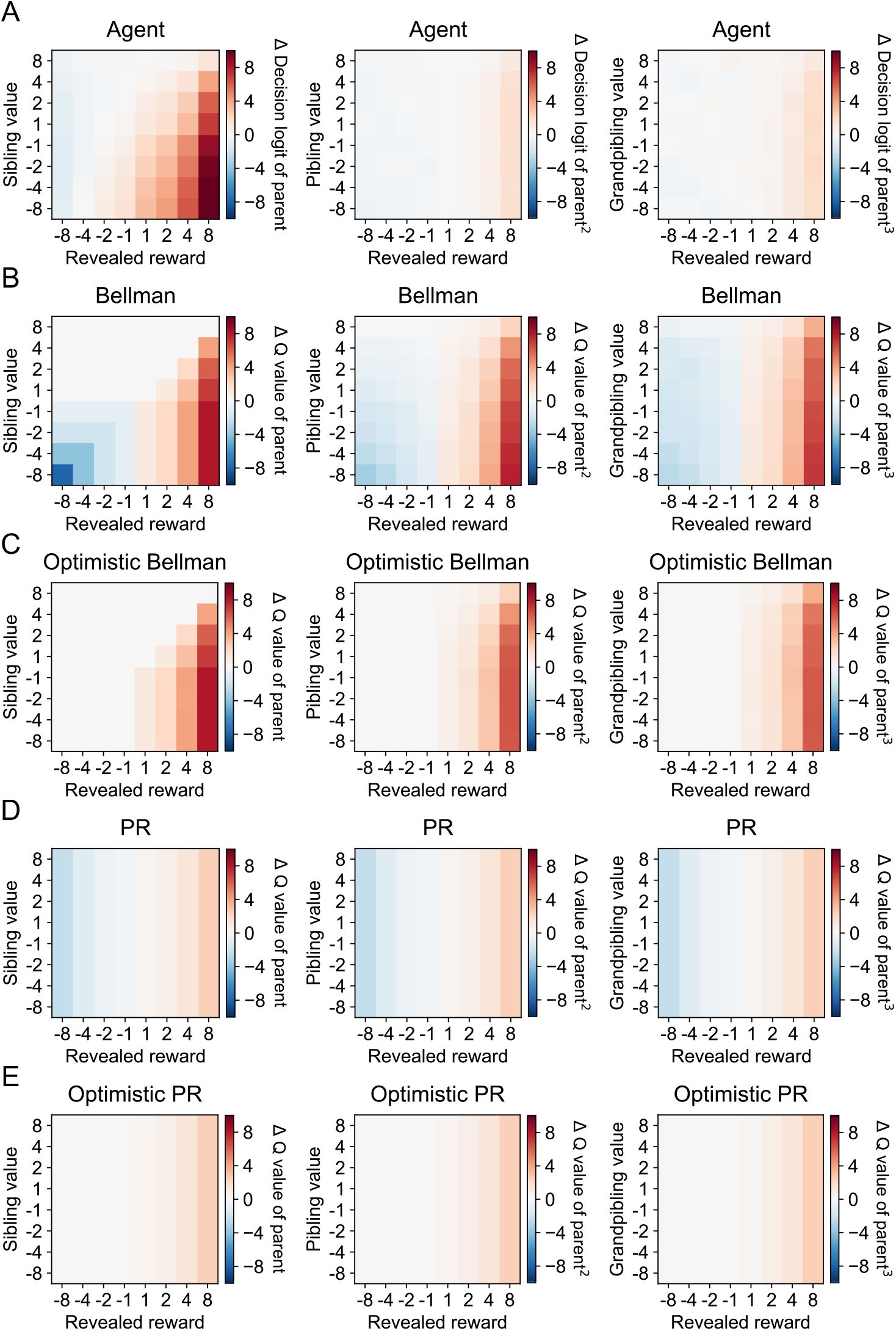
Comparison of backup strategies with baselines. We compared the agent’s backup strategies against four baselines: (i) Bellman backup: Q values are propagated to the parent by taking the maximum over children; (ii) optimistic Bellman backup: same as (i), but only positive Q values are propagated; (iii): predecessor representation: the agent tracks the identities of all state along the path to the current state, and each query increases of decreases the logits of all the nodes on that path in proportion to the reward at the queried state; and (iv) optimistic predecessor representation: same as (iii), but only positive rewards are propagated. For each scheme, we evaluated how the decision logit of the parent, grandparent, and great-grandparent changes as a function of the revealed reward and the value of the sibling, pibling (i.e., the parent’s sibling), and grandpibling (i.e., the grandparent’s sibling), respectively. We found that local backups are consistent with optimistic Bellman backups: the parent’s logit increases only when the revealed reward exceeds the sibling’s value. Non-local backups, however, appear less sensitive to the pibling’s or grandpibling’s value. They are more consistent with optimistic predecessor representation, in which propagation does not involve taking a maximum over children. We attribute this to memory constraints that limit the agent’s ability to perform such maximization over distal relatives.

**Figure S5:**
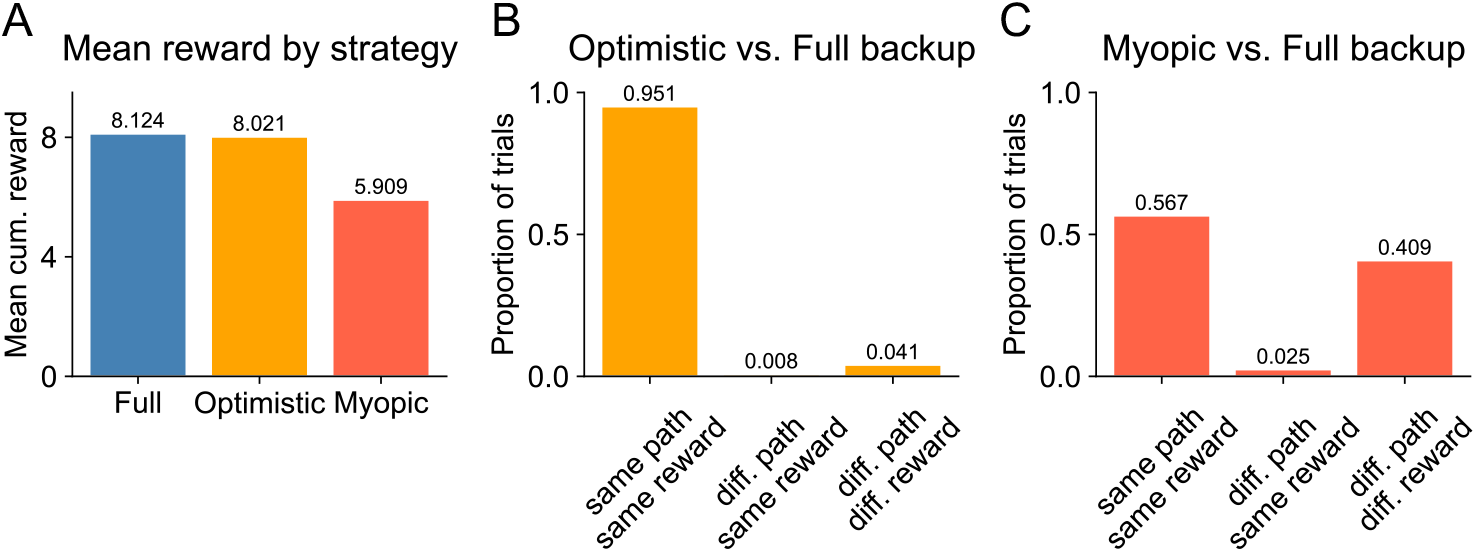
Optimistic backup is an effective heuristic. We evaluated whether optimistic backups achieve performance comparable to full Bellman backup. We compared three backup strategies: full Bellman backup, optimistic Bellman backup, and a myopic baseline (no backup), and found that optimistic backup closely matches full backup performance (A), with the two strategies predicting the same chosen path in 95% of trials (B). This confirms that optimistic backup is an effective heuristic and helps explain why the agent’s backup strategy shows an optimistic bias.

## Footnotes

1 Despite the name, “meta-reasoning” does not imply explicit reasoning about reasoning. Here, we define meta-reasoning as any computational process whose purpose is to control another computational process.

2 In a metalevel MDP, the effect of computation is encoded in the transition function, typically hand-specified to implement Bayesian inference over decision-relevant state. In our architecture, these effects correspond to the learned recurrent dynamics, *f*_*θ*_.

